# Genetically dissecting the electron transport chain of a soil bacterium reveals a generalizable mechanism for biological phenazine-1-carboxylic acid oxidation

**DOI:** 10.1101/2023.11.14.567096

**Authors:** Lev M.Z. Tsypin, Scott H. Saunders, Allen W. Chen, Dianne K. Newman

## Abstract

The capacity for bacterial extracellular electron transfer via secreted metabolites is widespread in natural, clinical, and industrial environments. Recently, we discovered biological oxidation of phenazine-1-carboxylic acid (PCA), the first example of biological regeneration of a naturally produced extracellular electron shuttle. However, it remained unclear how PCA oxidation was catalyzed. Here, we report the mechanism, which we uncovered by genetically perturbing the branched electron transport chain (ETC) of the soil isolate *Citrobacter portucalensis* MBL. Biological PCA oxidation is coupled to anaerobic respiration with nitrate, fumarate, dimethyl sulfoxide, or trimethylamine-N-oxide as terminal electron acceptors. Genetically inactivating the catalytic subunits for all redundant complexes for a given terminal electron acceptor abolishes PCA oxidation. In the absence of quinones, PCA can still donate electrons to certain terminal reductases, albeit much less efficiently. In *C. portucalensis* MBL, PCA oxidation is largely driven by flux through the ETC, which suggests a generalizable mechanism that may be employed by any anaerobically respiring bacterium with an accessible cytoplasmic membrane. This model is supported by analogous genetic experiments during nitrate respiration by *Pseudomonas aeruginosa*.

**Author summary:** Many bacteria have extremely flexible metabolisms, and we are only beginning to understand how they manifest in the environment. Our study focuses on the role of phenazine-1-carboxylic acid (PCA), a molecule that some bacteria synthesize and secrete into their surroundings. PCA is an “extracellular electron shuttle,” a molecule that readily transfers electrons between cells and oxidizing/reducing compounds or other cells. Until our investigation, the role of PCA electron-shuttling had only been studied in one direction: how it takes electrons away from cells, and the effect this has on their viability. Here we present a detailed account of the opposite process and its mechanism: what happens when PCA delivers electrons to cells? Our findings indicate that this previously underappreciated process is generalizable to any anaerobically respiring bacterium. Consequently, we expect that electron donation by PCA is widespread in environments where PCA is plentiful and oxygen is sparse, such as in some agricultural soils. The universality of the extracellular electron shuttle oxidation mechanism we describe for PCA suggests that it should also occur with similar small molecules, of which there are thousands, deepening the implication that this is a significant process in the environment and motivating further research into its consequences.

## Introduction

Phenazines are secreted secondary metabolites produced by diverse soil bacteria (1) that microbes use in various ways: from quorum sensing (2) to antimicrobial warfare (3–5) to energy conservation under anoxia (6). Each of these biological roles is connected to the ability of phenazines to accept and donate electrons (i.e., their redox activity), a process that has been studied for over 120 years. Bacterial phenazine reduction was first proposed in the nineteenth century as an indicator for the presence of enterics in water supplies (7). Several decades later, pyocyanin, one of the phenazines produced by *Pseudomonas aeruginosa*, was described as an “accessory respiratory pigment” that increased the rate of oxygen consumption by *Staphylococcus*, *Pneumococcus*, and erythrocytes by shuttling electrons from the cells to oxygen (8). Once it became apparent that phenazines can have cytotoxic effects, they were characterized as antimicrobial compounds that destructively abstract electrons from the transport chain (3). It was then discovered that reducing phenazines can greatly benefit *Pseudomonas aeruginosa* by 1) regulating gene expression during quorum sensing by oxidizing a transcription factor; 2) acting as alternative terminal electron acceptors to promote anoxic survival; and 3) facilitating nutrient acquisition (2,6,9–11). These reports paint a complex picture of the multifarious effects phenazines can have, but in each case, the conceptual model ends with the cell reducing the phenazine, which raises the question: how are phenazines recycled?

The first answer is: abiotically. Phenazines are broadly reactive molecules and can be oxidized by a variety of oxidants, including molecular oxygen and manganese or iron minerals. When oxygen serves as the electron acceptor, superoxide is produced, harming both phenazine producers and other cell types (12). In contrast, when iron minerals, to which phosphate is adsorbed, serve as the electron acceptor, ferrous iron and phosphate can be released, alleviating nutrient limitation (11,13,14). However, not all oxidants of higher redox potential (e.g., nitrate and nitrite) react quickly enough with phenazines to re-oxide them abiotically on biologically relevant timescales. Nonetheless, bacteria with versatile respiratory electron transport chains, such as the soil isolate *C. portucalensis* MBL, can catalyze biological phenazine oxidation under anoxic conditions with nitrate (15). This observation provides a second possible answer to the question of how phenazines are recycled: it stands to reason that in anoxic microenvironments where phenazine-reactive abiotic oxidants are limited, phenazine-reducing bacteria may benefit from the presence of phenazine-oxidizing bacteria. Furthermore, we hypothesize that phenazine oxidation itself might provide a survival advantage to phenazine-oxidizing cells under anaerobic conditions where electron donors are limited.

Towards testing these ideas, we used a systematic genetic approach to dissect how *C. portucalensis* oxidizes phenazines, focusing on potential catalysts within its electron transport chain (Figure 1A, Table 1). In addition to leading us to a generalizable mechanistic model for how biological PCA works, the development of a genetic system in this recently isolated soil organism exemplifies how the adaptation of existing tools can be used to rapidly gain new insights into microbial processes of environmental interest.

**Figure 1.**
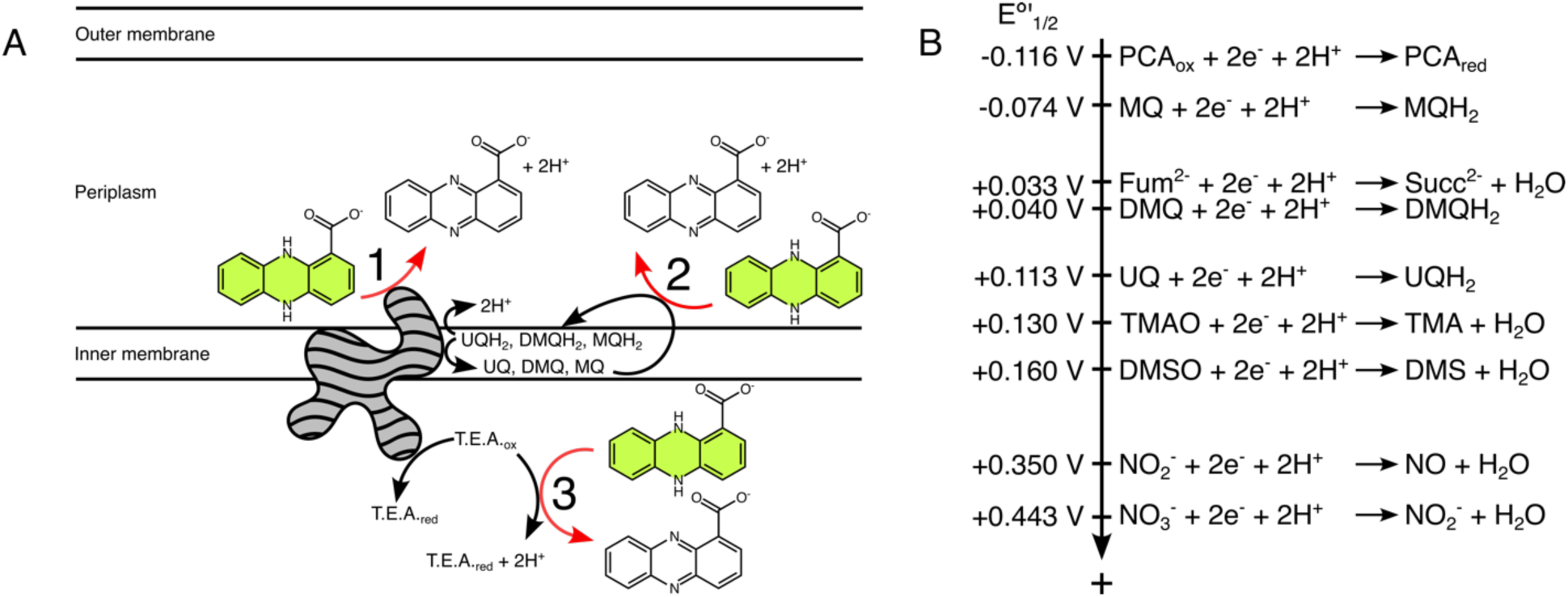
Models and thermodynamics of PCA oxidation. *(A) Potential models of PCA oxidation in Gram-negative bacteria capable of respiration.* In aqueous environments, PCA redox reactions are two-electron, two-proton processes. Reduced PCA is shown in green, reflecting its true color. PCA oxidation can theoretically be coupled to the respiratory electron transport chain in a couple ways: (Category 1) PCA may donate electrons to the terminal reductase (grey shape) for a respirable terminal electron acceptor (T.E.A.), thus contributing two protons to the periplasm; (Category 2) PCA may donate two electrons and two protons to quinones, thus regenerating the quinol pool. (Category 3) Alternatively, PCA directly reduce the terminal electron acceptor. This may happen externally to the cell, or PCA may enter the cytoplasm and react with the terminal electron acceptor independently of the electron transport chain, as depicted. In this illustration, the arrows may represent direct reactions or ones mediated by enzymes or other factors. Categories 1 and 2 represent scenarios that may be energetically beneficial for a respiring bacterial cell, whereas Category 3 may be detrimental. Category 3 would require transport of PCA across the inner membrane because its carboxylic acid moiety is negatively charged at circumneutral pH, and it cannot passively cross the membrane. For simplicity, this illustration does not show potential reactions with a periplasmic reductase, but the logic would remain the same, only with no involvement of the cytoplasmic space. *(B) Electron tower of relevant half-reactions.* Reactions are ordered by their relative standard midpoint potentials with more negative values on top and more positive ones on the bottom (not to scale). Thermodynamically favorable pairings comprise more positive half-reactions with more negative ones in reverse. The theoretical limit for energy that can be conserved from a pairing correlates with the magnitude of the different in half-reaction potentials. PCA: phenazine-1-carboxylic acid. MQ: menaquinone. DMQ: demethylmenaquinone. UQ: ubiquinone. Fum^2-^: fumarate. Succ^2-^: succinate. TMAO: trimethylamine-N-oxide. TMA: trimethylamine. DMSO: dimethyl sulfoxide. DMS: dimethyl sulfide. NO2-: nitrite. NO: nitric oxide. NO3-: nitrate.

**Table 1:** The categories of PCA oxidation reactions.

| Type | Description |
| --- | --- |
| Category 1 | PCA reduces the terminal reductase. |
| Category 2 | PCA reduces a quinone, thereby replenishing the quinol pool. |
| Category 3 | PCA reduces the terminal electron acceptor, either internally or externally to the cell (i.e., abiotic oxidation of PCA by the terminal electron acceptor). |

## Results

### A theoretical framework for understanding PCA oxidation metabolisms

In principle, many electron acceptors, both organic and inorganic, have the potential to oxidize PCA. Having previously found PCA oxidation to be coupled to nitrate reduction, we began our search for a general mechanism for PCA oxidation by performing a theoretical analysis of the reaction’s thermodynamics (Figure 1B). We considered the theoretical limit to the potential energy that could be conserved when PCA is oxidized by typical terminal electron acceptors used during anaerobic respiration. The thermodynamic favorability of these reactions is determined by their Gibbs free energy (ΔG). The ΔG of a given reaction (ΔG_r_) is related to the ΔG under standard conditions at pH (ΔG°′_r_):

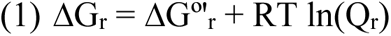

Q_r_ is the quotient of the activities of the products and reactants, T is the temperature in Kelvin, and R is the ideal gas constant (8.314 J K^-1^ mol ^-1^). ΔG°′_r_ can be derived from the midpoint potentials of reduction-oxidation (redox) reactions:

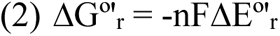

Here, n is the number of electrons exchanged, F is the Faraday constant (96,485 J mol^-1^ V ^-1^), and ΔE°′_r_ is the difference in the standard potential of the reactants. In the case of nitrate and phenazine-1-carboxylic acid (PCA), the analysis gives the following result:

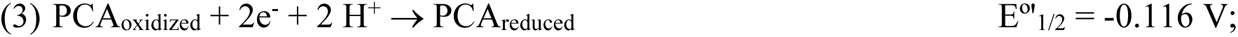

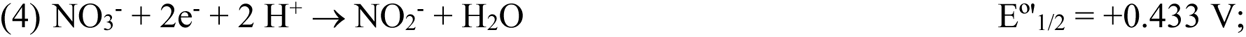

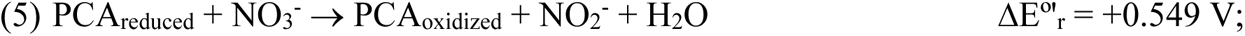

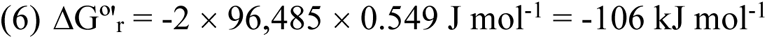

Thus, under standard conditions at pH 7, the coupling of PCA oxidation to nitrate reduction gives a negative Gibbs free energy, meaning that it would be thermodynamically favorable. However, while we can control the pressure, temperature, and pH under laboratory conditions, there is no reason for the concentration conditions to be standard, and so it is important to consider at which ratio Q_r_ the reaction will become *unfavorable*. This can be derived as follows:

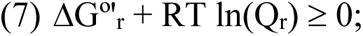

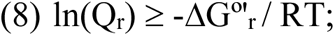

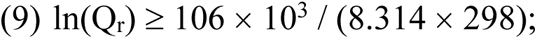

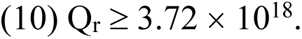

The coupling of PCA oxidation to nitrate reduction becomes unfavorable when the ratio of the activities of the product and reactant is over 18 orders of magnitude. In effect, this reaction will be favorable until it reaches completion, which matches our observation with *C. portucalensis* MBL, our model organism for studying PCA oxidation (15,16).

The same theoretical analysis applies to the other alternative terminal electron acceptors: fumarate (E°′_1/2_ = +0.033 V), DMSO (E°′_1/2_ = +0.160 V), TMAO (E°′_1/2_ = +0.130 V), and nitrite (E°′_1/2_ = +0.350 V) (Table 2) (17,18). During respiration, electrons travel from cytoplasmic reducing equivalents, such as NADH (E°′_1/2_ = -0.320 V) (17), to the terminal electron acceptors via flux through the quinone/quinol pool in the cytoplasmic membrane. Conceivably, during their biologically catalyzed oxidation, phenazines may also donate their electrons to quinones. *C. portucalensis* MBL has the genes to synthesize three quinones: ubiquinone (UQ; E°′_1/2_ = +0.113 V), menaquinone (MQ; E°′_1/2_ = -0.074 V), and demethylmenaquinone (DMQ; E°′_1/2_ = +0.040 V) (Figure 2 and Supplementary Figure 2.1). Based on their respective midpoint potentials, each of these would be expected to accept electrons from PCA (Figure 1B and Table 2) (17,18). Thus, it is feasible for bacteria to use these reactions during their metabolism.

**Figure 2.**
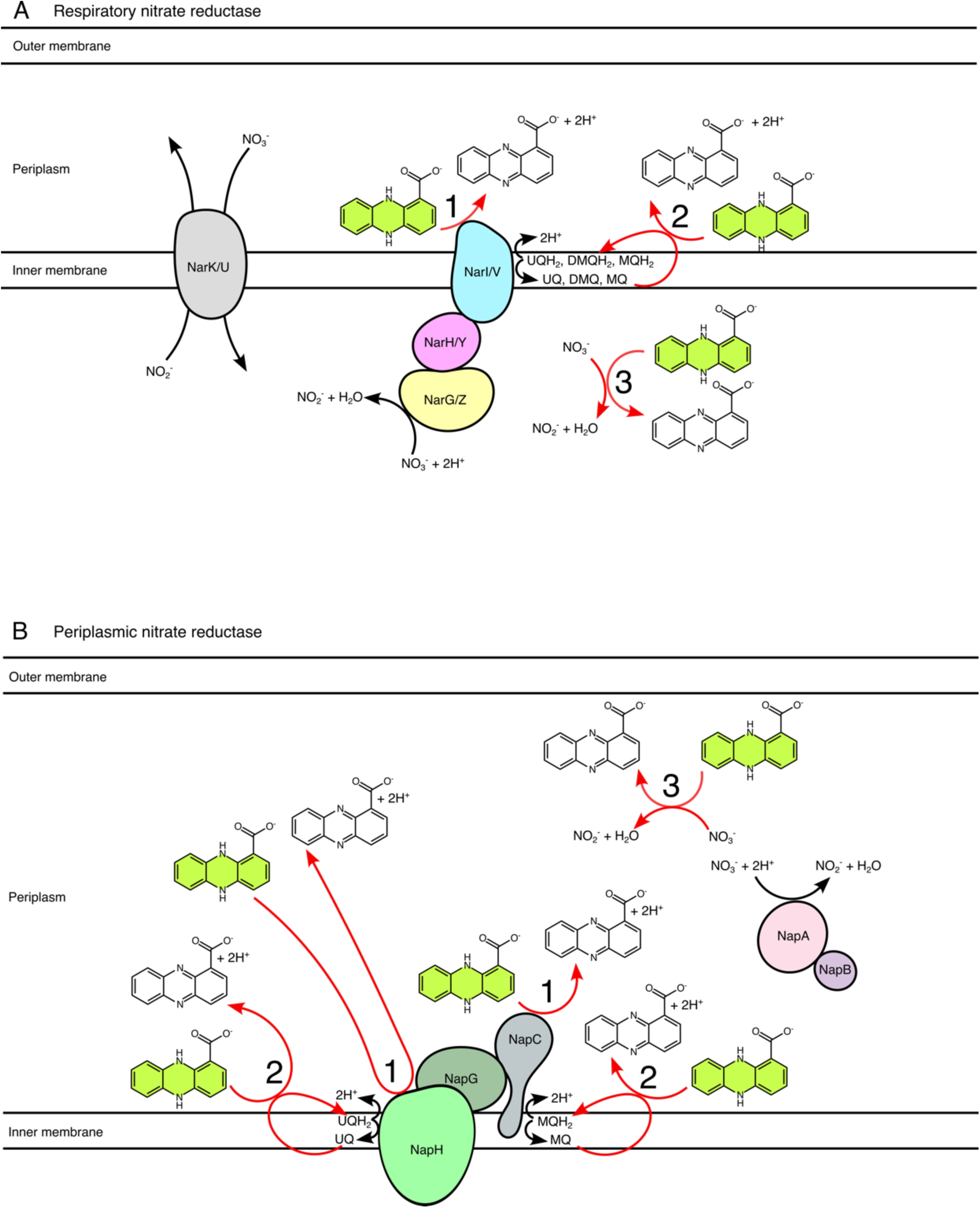
Explicit conceptual model of PCA oxidation during anaerobic nitrate respiration. To interpret PCA oxidation phenotypes during anaerobic nitrate respiration, it is necessary to keep in mind the distinct electron pathways in the respiratory (A) and periplasmic (B) nitrate reductase complexes. These models are made assuming complete analogy to the arrangement of these proteins in *E. coli* (18,20,21). The paths of the electrons along the reductase complexes are not shown for simplicity, but they flow from the quinols to the nitrate in each case. The categories of PCA interactions are numbered according to the scheme in Figure 1A. In the case of the respiratory nitrate reductase complexes, there are two redundant homologs in *C. portucalensis*: NarGHI and NarZYV (16). NarI/V can accept electrons from all three types of quinones (18). For the periplasmic nitrate reductase, there are two distinct quinone interaction sites (NapH for ubiquinone and NapC for menaquinone); notably, demethylmenaquinone does not appear to play a role in periplasmic nitrate reductase activity (21). The NapAB complex is soluble in the periplasmic space, and the electrons from the NapHGC complex are ferried to NapA by NapB, which is a cytochrome c-type protein (20). Category 1 PCA interactions are depicted as occurring at quinol-oxidizing subunits of the protein complexes (NarI, NarV, NapH, and NapC) to illustrate the hypothesis that a reduced PCA molecule may replace a quinol. Auxiliary and chaperone proteins that are members of the nitrate reductase operons and are involved in complex formation but not activity (NarJ, NarW, NapF, and NapD) are not shown. Note: *P. aeruginosa* possesses only one set of homologs for the respiratory nitrate reductase (NarGHI) and the periplasmic nitrate reductase.

**Table 2.** Thermodynamics of PCA oxidation coupled to anaerobic terminal electron acceptor reduction.

| Electron acceptor | $\Delta G^{\circ}_r$ (kJ mol<sup>-1</sup>) | Unfavorable boundary |
| --- | --- | --- |
| Nitrate | -106 | $Q_r \geq 3.72 \times 10^{18}$ |
| Nitrite | -89.9 | $Q_r \geq 5.79 \times 10^{15}$ |
| DMSO | -53.3 | $Q_r \geq 2.17 \times 10^9$ |
| TMAO | -47.5 | $Q_r \geq 2.09 \times 10^8$ |
| Ubiquinone (UQ) | -44.2 | $Q_r \geq 5.57 \times 10^7$ |
| Demethylmenaquinone (DMQ) | -30.1 | $Q_r \geq 1.89 \times 10^5$ |
| Fumarate | -28.8 | $Q_r \geq 1.10 \times 10^5$ |
| Menaquinone (MQ) | -8.10 | $Q_r \geq 26.3$ |

Different terminal reductases take electrons from different quinones, as determined by thermodynamic and kinetic constraints. For instance, nitrate has a sufficiently positive midpoint potential to favorably oxidize any of the quinones, while fumarate can only drive menaquinone oxidation to completion (Figure 1B). These redox relationships enable a conceptual model for interpreting PCA oxidation phenotypes: For instance, if ubiquinone is necessary for PCA oxidation, fumarate would not stimulate the process because its midpoint potential is such that it cannot oxidize ubiquinol (Figure 1B). The fact that PCA is abiotically oxidized by certain compounds leads to the hypothesis that biological PCA oxidation does not require specialized enzymes. To test this, we can distinguish whether PCA donates electrons to the terminal reductases via the quinol pool and, if it does, which of the quinones are at play. The hypothesis that quinones mediate phenazine oxidation in the bacterial electron transport chain is supported be the observation that ubiquinone-1 rapidly oxidizes the phenazines pyocyanin (PYO), phenazine carboxamide (PCN), and PCA in aqueous solution (19).

Following the schemes in Figures 1 and 2, we set out to employ genetic perturbations to test which categories of PCA oxidation can proceed (Table 1). For example, after ruling out Category 3 reactions with abiotic controls, we knocked out the available quinones and asked whether Category 1 and 2 reactions occur: if no quinones were present and oxidation still proceeded, then Category 1 PCA oxidation played a role; if the rate of Category 1 reactions could not explain the wildtype PCA oxidation rate alone, then Category 2 reactions were co-occurring. Likewise, knocking out the terminal reductases themselves was a critical test: if we could not fully abolish PCA oxidation to the level of the abiotic control by removing the terminal reductase(s), then there was a PCA oxidation mechanism that we missed. In the following results, we explore this genetic logic for understanding anaerobic PCA oxidation by *C. portucalensis* MBL for its four alternative respiratory TEAs (nitrate, fumarate, DMSO, and TMAO). For nitrate-driven PCA oxidation, we also provide a validation against *Pseudomonas aeruginosa*, which lacks the ability to respire the other TEAs and has a reduced complement of quinones (Supplementary Figure 2.1).

### Inactivation of individual nitrate reductases yields mild PCA oxidation defects

To investigate the mechanisms and dynamics of PCA oxidation by *C. portucalensis* MBL, we adapted *Escherichia coli* genetic engineering protocols. Given that *C. portucalensis* harbors three functionally redundant nitrate reductase complexes (whose catalytic subunits comprise NapA, NarG or NarZ), we hypothesized that inactivation of any one of these enzymes would be insufficient to abolish PCA-oxidation activity. Accordingly, we sought to develop an efficient method that would permit the combinatorial disruption of genes encoding these enzymes.

To start, we compared the phenotypes of strains in which we perturbed nitrate reductases by two different methods: λRed recombination or oligo-mediated recombineering to make whole operon deletions or targeted translational knockouts of catalytic subunits, respectively (26,27). In the λRed approach, the operons were replaced by a kanamycin resistance cassette by homologous recombination using the λRed recombinase expressed on a transient plasmid (26). In the oligo-mediated recombineering approach, we replaced three subsequent codons with the three different stop codons (TAA, TGA, and TAG, not necessarily in that order) in the first half of the coding sequence of the catalytic subunit of the given gene(s). Figure 2 shows the three nitrate reductases we mutagenized, as well as their predicted orientations in the ETC. According to this model, deleting the entire operon or knocking out just the catalytic subunit should have the same effect: all three categories of PCA oxidation reactions would be abolished.

As expected for redundant systems, disruption of any single nitrate reductase by either genetic engineering method could reduce, but not eliminate, PCA oxidation. We observed no difference between the *narZ* translational knockout (*narZ-tlKO)* and the operon deletion (*ΔnarZYWV*) in the rate and dynamics of PCA oxidation (Figure 3, left). Comparing *narZ-tlKO* and *ΔnarUZYWV*, we found that *ΔnarUZYWV* had a greater delay before PCA oxidation commenced, but the maximum rate of PCA oxidation was the same as for *narZ-tlKO* and *ΔnarZYWV* (Figure 3, left). All three mutants oxidized PCA later and more slowly than the wildtype (Figure 3, left). The lag in oxidation by the *ΔnarUZYWV* strain compared to the Δ*narZYWV* and *narZ-tlKO* strains may reflect the fact that *narU* is an inner membrane nitrate-nitrite antiporter (28): its absence may delay nitrate entrance to the cytosol where it can encounter NarZ or NarG (Figure 2). As observed for the *narZ* mutants, there was no difference between the *narG-tlKO* and *ΔnarGHJI* PCA oxidation phenotypes, and there was only mild loss of PCA reduction compared to the wildtype (Figure 3, middle). The *napA-tlKO* strain appeared to have a more severe PCA oxidation defect than *ΔnapFDAGHBC* versus the wildtype, yet loss of PCA oxidation by these strains still only minimally slowed down PCA oxidation (Figure 3, right). For the rest of this study, we used oligo recombineering rather than λRed recombination to generate strains with multiple gene disruptions.

**Figure 3.**
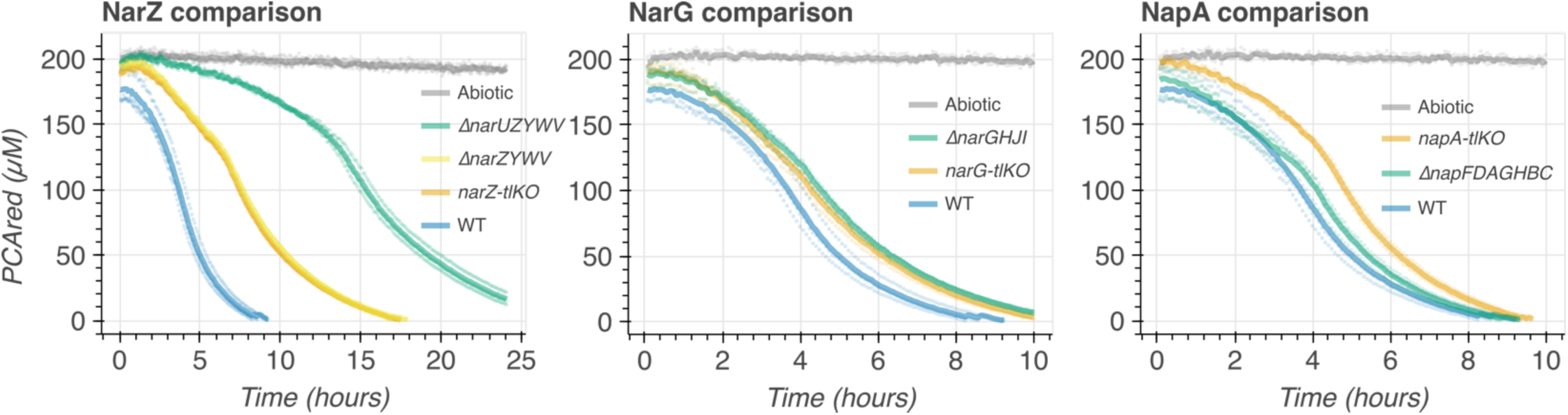
Comparison of operon deletions and catalytic subunit translational knockouts for terminal nitrate reductases. Each graph shows the oxidation of PCA over time (measured by the decay of PCAred, which is fluorescent) with the abiotic control (nitrate plus reduced PCA in reaction medium without cells) in grey. Left, NarZ comparison: the deletion (Δ) and translational knockout (tlKO) strains all have a PCA oxidation deficit relative to the wildtype (WT, blue). The most severe phenotype is in the *ΔnarUZYWV* strain (green), and the *ΔnarZYWV* and *narZ*-tlKO (yellow and orange, respectively) phenotypes are undistinguishable. Middle, NarG comparison: the deletion (green) and translational knockout (orange) strains have the same slight PCA oxidation deficit relative to the wildtype control (WT, blue). Right, NapA comparisons: the translational knockout (orange) has a more severe PCA oxidation deficit that the deletion strain (green), relative to the wildtype control (blue). Each thick line corresponds to the mean of three biological replicates plotted in semitransparent circles.

### PCA oxidation dynamics with nitrate depend on which nitrate reductases are present

Given the presence of three nitrate reductases in the genome, we speculated that our ability to alter PCA oxidation dynamics via their deletion, alone or in combination, would depend on their expression under different growth conditions. Accordingly, we compared nitrate-driven PCA oxidation dynamics after two different pregrowth conditions in lysogeny broth (LB): slanted shaking overnight tubes and standing parafilm-sealed overnight tubes. These two conditions permitted fully aerated (oxic) and hypoxic growth, respectively. Depending on which cultivation condition was used, distinct nitrate reductases dominantly contributed to PCA oxidation (Figure 4). The PCA oxidation dynamics and phenotypes can generally be summarized by the time it takes to reach a threshold concentration of reduced PCA (e.g., Figure 4A) and the maximal PCA oxidation rate (e.g., Figure 4B), but this can obscure some more nuanced phenotypes, such as the biphasic nature of many of the PCA oxidation curves (e.g., Fig 3 *narZ-tlKO*). While the fit and oxidation metrics depend on the parametrization of the model, we were able to identify a conservative parametrization that robustly represented all tested strains (Supplementary Figure 4.1).

**Figure 4.**
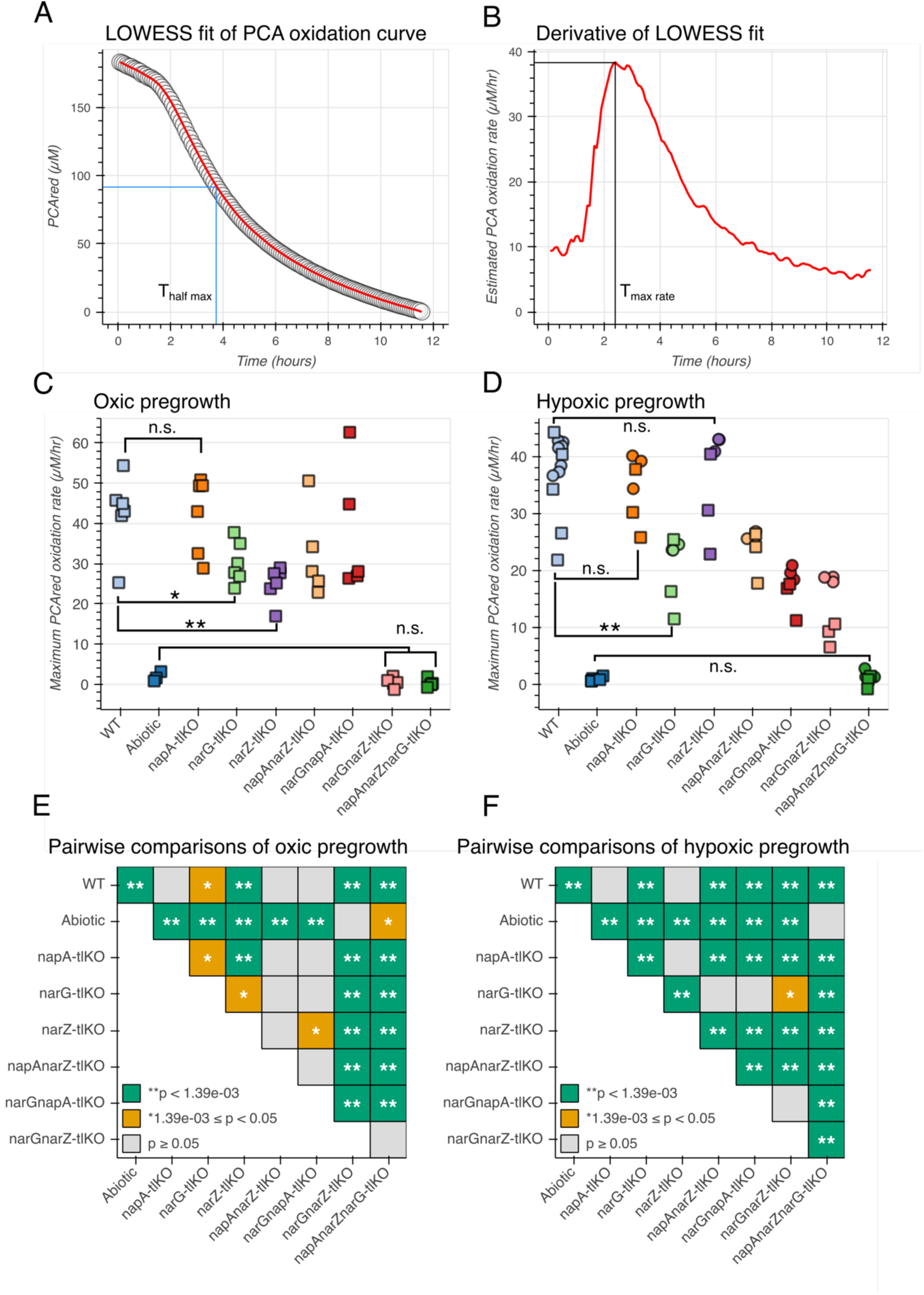
PCA oxidation dynamics of nitrate reductase knockouts, comparing shaking (oxic) and standing (hypoxic) overnight pregrowth. (A-B) A demonstration of the analysis pipeline. (A) We used a locally weighted scatterplot smoothing (LOWESS) algorithm to fit a curve to our empirical data, here showing an example of one of the biological replicates for wildtype *C. portucalensis* MBL oxidizing PCA with nitrate after hypoxic overnight pregrowth. The data are in semitransparent circles and the LOWESS fit is the red line. The fitting was parametrized to use 5% of the data in its sliding window. This fit allowed us to determine the time it took the cultures to oxidize half of the provided PCA (Thalf max, or any arbitrary threshold) and the derivative, or rate, of PCA oxidation. (B) This derivative allowed us to estimate the maximum rate of PCA oxidation and the time at which it occurred. (C) The maximum PCA oxidation rate in the presence *C. portucalensis* MBL strains of each nitrate reductase genotype, as well as the abiotic condition, after oxic pregrowth. Squares represent the means of technical triplicates. (D) Similarly, the maximum PCA oxidation rate of the different strains and controls after hypoxic overnight pregrowth. In both (C) and (D), most biologically illuminating comparisons are between the mutants and the wildtype and abiotic controls. This allows us to observe that in the oxic pregrowth condition, NapA does not contribute to PCA oxidation at all, as evidenced by the fact that both the *narGnarZ* and *napAnarZnarG* strains are indistinguishable from the abiotic control. Meanwhile, only the *narZ* single knockout has a phenotype relative to the control. In contrast, each of the three nitrate reductases contributes to PCA oxidation after hypoxic pregrowth (only the triple knockout is indistinguishable from the control), but only *narG* alone has a phenotype relative to the wildtype. n.s. stands for “not significant”. (E-F) Pairwise statistical tests for all the strains and controls tested after both oxic (E) and hypoxic (F) pregrowth conditions using the null hypothesis that there is no difference between the mean maximum oxidation rates of two given genotypes (or the abiotic control). Given 36 comparisons, the Bonferroni-corrected p-value threshold for significance is p < 0.00139. See the methods for how the statistical significance testing was performed.

After oxic pregrowth (Figure 4C), the *narZ* knockout had the most severe phenotype of the single mutants, and eliminating both *narG* and *narZ* was sufficient to abolish PCA oxidation, implying that under this condition *napA* is irrelevant (Figure 4E). The time it took for the different strains to oxidize 50% of the provided reduced PCA was correlated with the average maximum PCA oxidation rate, and the *narZ* single knockout was again the slowest among the other single mutants (Supplementary Figure 4.2A and C). The abiotic, *narGnarZ* double knockout, and *napAnarZnarG* triple knockout conditions never reached this 50% threshold (Supplementary Figure 4.2A). Relative to the wildtype, only the *narZ* single knockout strain had a significantly different time to 50% oxidation (Supplementary Figure 4.2A).

Cells pre-grown in stationary cultures provided a subtle but important contrast to the shaking pregrowth results (Figure 4D and F). Here, each of the double knockouts had a detectable PCA oxidation rate, indicating that any of the three nitrate reductases can drive the oxidation (Figure 4D and F). Moreover, rather than *narZ* above, the greatest loss of oxidation in a single mutant background came from the *narG* (Figure 4D). Once again, the time to 50% oxidation was correlated to the maximum oxidation rate (Supplementary Figure 4.2B), but in this condition, the *narZ* single mutant did not delay reaching this threshold relative to the wildtype activity (Supplementary Figure 4.2B and D). These results imply that the pregrowth condition determines the relative presence and activity of the different nitrate reductases prior to the oxidation assay, leading to different phenotypes for a given strain. Because pre-growth in stationary cultures allowed us to observe the contributions of each of the three nitrate reductases, we employed this condition for the rest of the genetic experiments, including complementation assays (Supplementary Figure 4.3). When the nitrate reductases were individually overexpressed in the triple knockout background during overnight stationary pre-growth (Supplementary Figure 4.3A), only the *narZ* overexpression strain had a statistically significant rescue of PCA oxidation (Supplementary Figure 4.3B). The rescue effect was small, and the *napA* and *narG* overexpression strains did shift toward higher oxidation rates, so this is likely due to the overexpression system being unoptimized for *C. portucalensis* MBL.

### Loss of quinones in *C. portucalensis* MBL significantly disrupts PCA oxidation, yet mild oxidation is still achieved by nitrate reductases in their absence

Nitrate-driven PCA oxidation by *C. portucalensis* is fully abolished only when all three terminal nitrate reductases are knocked out; any one remaining reductase enables PCA oxidation (Figure 4D). This observation raises the question whether the nitrate reductases oxidize PCA directly (Figure 2 and Table 1, Category 1 reactions) or if PCA contributes electrons to an upstream pool, such as the quinols (Figure 2 and Table 1, Category 2 reactions). PCA oxidation was largely lost when the biosynthesis of all three quinones (ubiquinone, menaquinone, and demethylmenaquinone) was disrupted in the *menAubiC*-tlKO strain, indicating that Category 2 PCA oxidation reactions are dominant (Figure 5A). However, the PCA oxidation rate in this genetic background did not drop to abiotic levels, indicating that cellular nitrate reduction can drive PCA oxidation at a low rate without quinones as intermediaries (Figure 5A). In other words, Category 1 PCA oxidation reactions also occur. The loss of ubiquinones alone in the *ubiC-tlKO* background had no effect on PCA oxidation, but the loss of (demethyl)menaquinones in the *menA*-*tlKO* strain did have an effect (Figure 5A and B). In the quinone-null background, PCA oxidation persists with any one of the nitrate reductases (Figure 5A), as evidenced by the positive oxidation rate in the *menAubiCnapAnarZ-*, *menAubiCnarGnapA-,* and *menAubiCnarGnarZ-tlKO* strains. Interestingly, the strains with no quinones and only intact *NarG* or *NapA* had faster PCA oxidation rates than the *menAubiC* strain on its own (Figure 5A and B). The co-occurrence of Category 1 and 2 reactions implies that PCA oxidation may be plastic: it does not necessarily depend on an specifically evolved enzyme or pathway to proceed.

**Figure 5.**
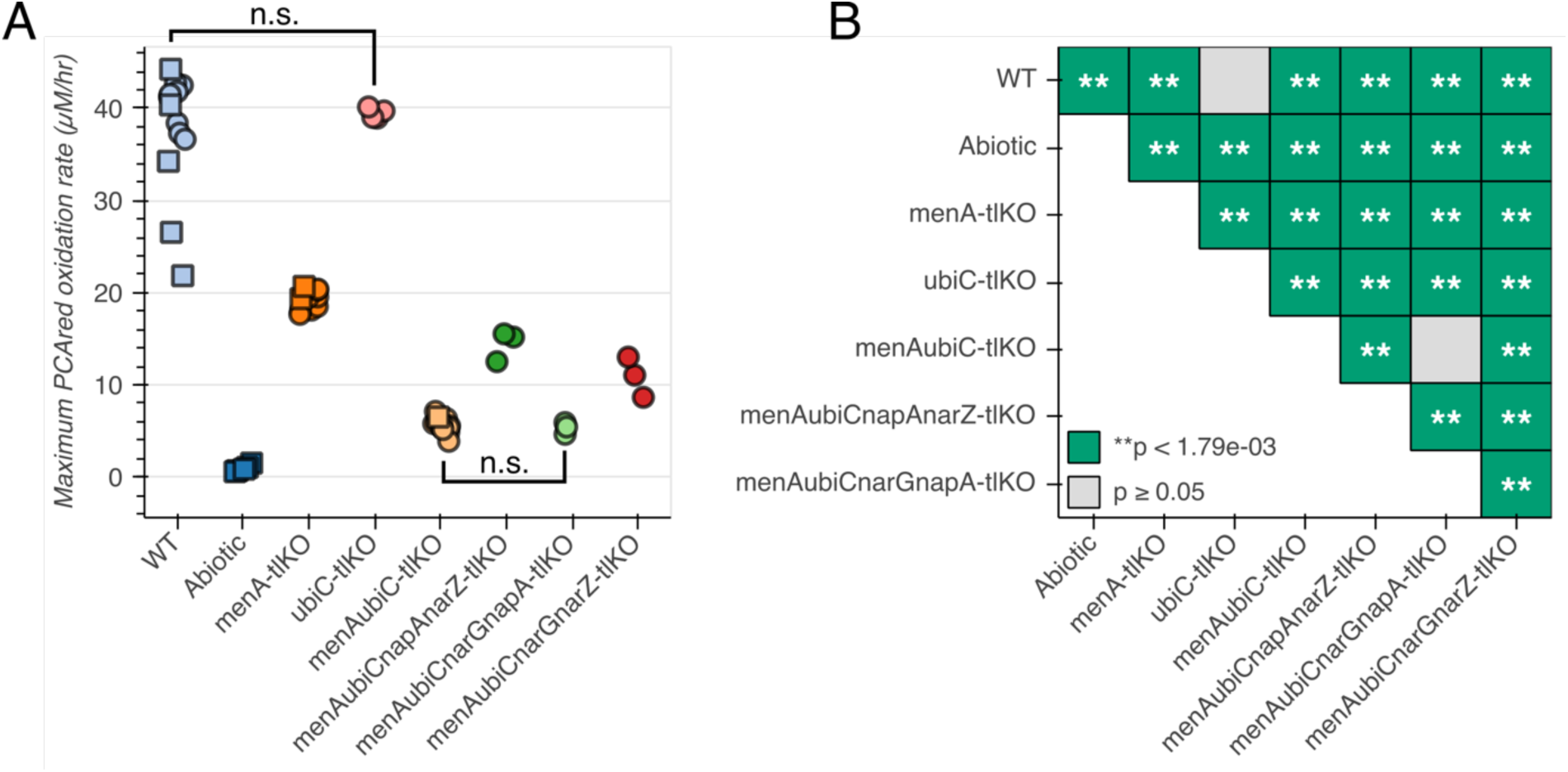
Nitrate-driven PCA oxidation rates in quinone knockout backgrounds, including single nitrate reductase strains. (A) Maximum PCA oxidation rates. Squares represent the means of technical triplicates and circles represent independent biological replicates. (B) Pairwise comparisons between differences of mean maximum PCA oxidation rates for the strains in (A). Given 28 comparisons, the Bonferroni-corrected threshold for significance is p < 0.00179.

We were unable to genetically complement the loss of quinones during nitrate-driven PCA oxidation (Supplementary Figure 5.1A and B). Curiously, overexpressing *menA* exacerbated the *menAubiC-tlKO* phenotype (Supplementary Figure 5.1A). Because each nitrate reductase retains some PCA oxidation activity even in the absence of quinones (Figure 5A), it may be the case that exogenously expressing a low level of quinones is not enough to give a signal over the independent nitrate reductase activity. However, our greater degree of success with complementing *menA* for the other terminal electron acceptors indicates that the system works in principle (see the alternative TEA section, below).

### Pseudomonas aeruginosa PA14 replicates nitrate reductase knockout phenotypes from C. portucalensis MBL

Given the apparent generalizability of PCA oxidation to the three nitrate reductases in *C. portucalensis* MBL and the observation that other bacteria perform the same metabolism (15,29), we assessed whether *P. aeruginosa* PA14 nitrate reductase mutants conform to the same mechanistic model. *P. aeruginosa* is particularly relevant as a point of comparison because it biosynthesizes PCA, has been studied extensively as a PCA reducer, and, like *C. portucalensis* MBL, also has both respiratory and periplasmic nitrate reductases (albeit only one homolog of the respiratory nitrate reductase complex, NarGHI) (Figure 2) (6,19,30). We compared wildtype, *ΔnarG*, and *ΔnapAB* strains of *P. aeruginosa* PA14 after pregrowth in shaking and standing conditions, as we had done for *C. portucalensis* MBL. We observed that in the shaking pregrowth condition, the *ΔnarG* strain had no phenotype versus the wildtype, while the *ΔnapAB* strain had a severe, though incomplete, PCA oxidation defect (Figure 6, left). In contrast, after either of the standing pregrowth treatments, the *ΔnapAB* strain did not substantially differ from the wildtype, while the Δ*narG* strain had the severe, incomplete loss of PCA oxidation (Figure 6, right). This pattern of distinct nitrate reductases dominating PCA oxidation activity depending on the pre-growth condition corresponds to the fact that *napA* and *narG* are regulated by distinct systems: RpoS and Anr, respectively (31,32). These phenotypes are analogous to the difference we observed between *C. portucalensis* MBL *narG* and *narZ* translational knockouts depending on pre-growth condition (Figure 4C and D), which implies that the PCA oxidation mechanism is not specific to a given organism or enzyme but rather the architecture of the electron transport chain.

**Figure 6.**
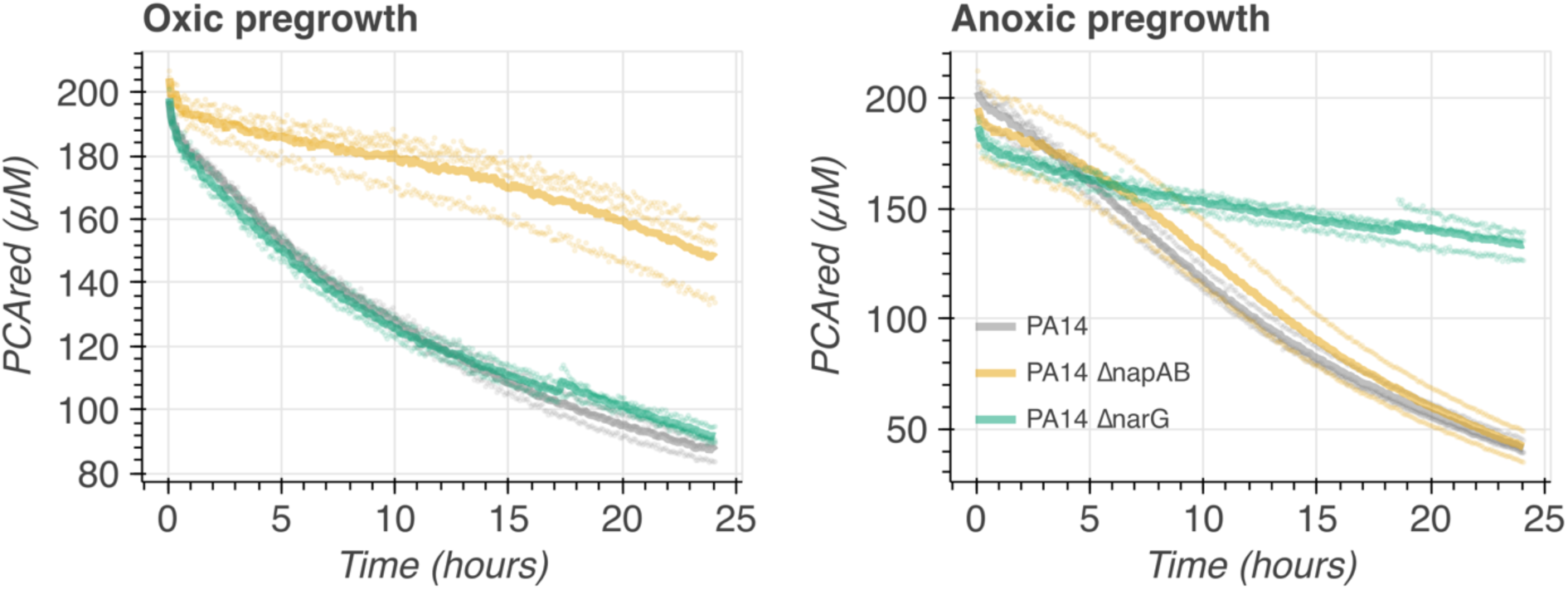
Pre-growth dependent phenotypes for P. aeruginosa nitrate reductase mutants. Left: When cultures were pre-grown in slanted shaking tubes, providing the cultures ample oxygen, only the *ΔnapAB* strain (orange) had a PCA oxidation deficit. Right: Pre-growing the cultures in standing tubes without nitrate supplementation also leads to the Δ*narG* strain (green) to have the PCA oxidation deficit; the wildtype (grey) and *ΔnapAB* (orange) strains are within error of each other. Each thick line corresponds to the mean of three biological replicates that are plotted in semitransparent circles. The medium for this experiment was 84% LB and 16% basal PCA oxidation assay medium (see materials and methods). Note: due to its relative inability to grow under hypoxia in LB, it was not feasible to grow a *nap/nar* double knockout P. aeruginosa strain to compare to the *C. portucalensis* MBL full nitrate reductase knockout. Similarly, we could not generate any quinone knockouts for *P. aeruginosa* because it encodes the biosynthesis of only ubiquinone (24).

### Alternative TEAs drive PCA oxidation analogously to nitrate

Having found a generalizable mechanism for nitrate-driven PCA oxidation in *C. portucalensis* MBL, we tested whether it extended to the other anaerobic respiratory metabolisms. We found that fumarate (terminal reductase catalytic subunit FrdA), DMSO (terminal reductase catalytic subunit DmsA), and TMAO (terminal reductase catalytic subunit TorA) can all drive PCA oxidation by *C. portucalensis* MBL (Figure 7). Analogously to nitrate-driven PCA oxidation, there was no abiotic reaction between reduced PCA and these alternative terminal electron acceptors. Knocking out the terminal reductase was sufficient to abolish fumarate-driven PCA oxidation (Figure 7A). In fact, the *frdA* knockout strain quenched residual oxygen, thus lowering the apparent oxidation rate beyond that of the abiotic control (Figure 7A and B). The *menA* knockout strain, lacking (demethyl-)menaquinones lost some PCA oxidation activity, but the *ubiC* knockout strain was indistinguishable from the wildtype (Figure 7A and B). Unlike in the case of nitrate-driven PCA oxidation (Figure 5A and B), the *menAubiC* knockout strain did not have a more severe phenotype than the *menA* knockout alone, which corresponds to the standard model that the fumarate reductase does not engage ubiquinone (18). It is also consistent that MenA complementation strain rescues some activity, and the UbiC complementation strain does not. In the complementation assays, the *frdA* overexpression vector was able to rescue roughly half of the wildtype activity over the *frdA* knockout background (Figure 7A and B, *frdA-tlKO* vs. *frdA* pFE21-FrdA), and the *menA* overexpression vector rescued PCA oxidation rates entirely (Figure 7A and B, WT vs. *menAubiC-tlKO* vs. *menAubiC* pFE21-MenA). The *ubiC* overexpression vector gave no effect (Figure 7A and B, *menAubiC-tlKO* vs. *menAubiC* pFE21-UbiC). Overall, the results here are analogous to what we observed while exploring nitrate-driven PCA oxidation: the abiotic control rules out Category 3 PCA oxidation, the *frdA* knockout abolishes PCA oxidation entirely (just as the *napAnarZnarG* triple knockout did for nitrate), and the full loss of quinones retains partial PCA oxidation activity, which indicates that both Category 1 and 2 PCA oxidation reactions play a role.

**Figure 7.**
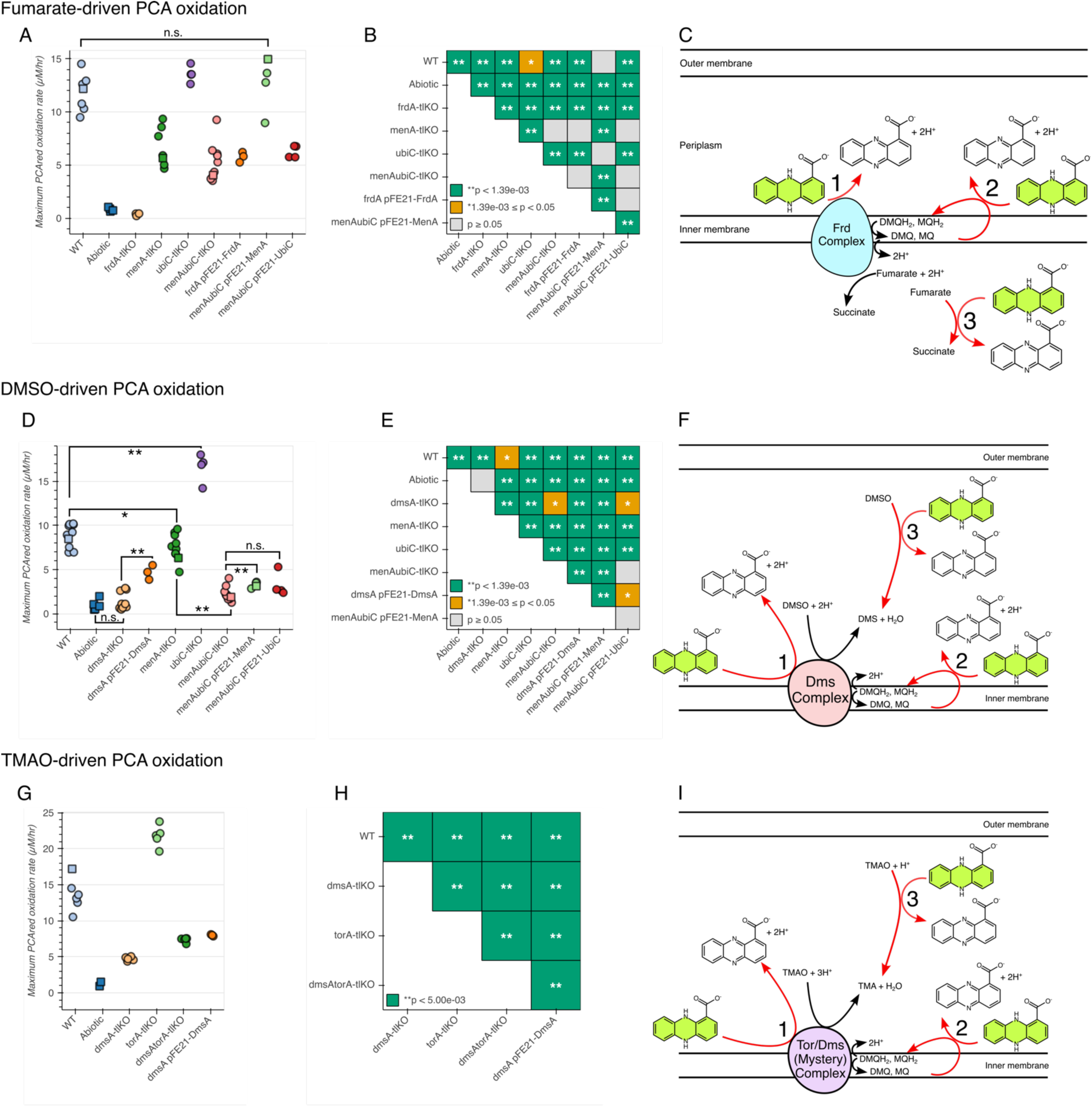
PCA oxidation driven by alterative TEAs in their respective reductase and quinone knockout strains, as compared to corresponding complementations. (A-C) Fumarate-driven PCA oxidation. (A) Maximum PCA oxidation rates. Squares represent the means of technical triplicates and circles represent independent biological replicates. (B) Pairwise comparisons between differences of mean maximum PCA oxidation rates for the strains in (A). Given 36 comparisons, the Bonferroni-corrected threshold for significance is p < 0.00139. (C) The ETC architecture for fumarate respiration, modeled on the *E. coli* literature (35,36). (D-F) DMSO-driven PCA oxidation. (D) Maximum PCA oxidation rates. Squares represent the means of technical triplicates and circles represent independent biological replicates. (E) Pairwise comparisons between differences of mean maximum PCA oxidation rates for the strains in (D). Given 21 comparisons, the Bonferroni-corrected threshold for significance is p < 0.00238. (F) The ETC architecture for DMSO respiration, modeled on the *E. coli* literature (37,38). (G-I) TMAO-driven PCA oxidation. (G) Maximum PCA oxidation rates. Squares represent the means of technical triplicates and circles represent independent biological replicates. (H) Pairwise comparisons between differences of mean maximum PCA oxidation rates for the strains in (G). Given 10 comparisons, the Bonferroni-corrected threshold for significance is p < 0.005. (I) The ETC architecture for TMAO respiration, modeled on the *E. coli* literature (34,39).

DMSO-driven PCA oxidation followed largely the same patterns as for nitrate and fumarate, with the surprising exception of ubiquinone phenotypes. There was no abiotic activity with DMSO, ruling out Category 3 PCA oxidation (Figure 7D and E). The *dmsA* knockout lost all activity, which was partially complemented by the DmsA overexpression strain (Figure 7D and E). The *menA* knockout did not have a strong phenotype, but the *ubiC* knockout had a substantially faster PCA oxidation than the wildtype (Figure 7D and E). However, the *menAubiC* double knockout had a severe loss of PCA oxidation, though still measurable above the abiotic control (Figure 7D and E). The MenA complementation strain slightly rescued PCA oxidation relative to the *menAubiC* background, but the UbiC complementation strain had no rescue. These results again support the roles of both Category 1 and 2 PCA oxidation reactions, but our model does not explain why loss of ubiquinones alone would increase PCA oxidation, while loss of ubiquinones on top of (demethyl-)menaquinones would decrease it (Figure 7F).

Our exploration of TMAO-driven PCA oxidation is less complete because we were not able to fully abolish PCA oxidation by knocking out the catalytic subunits of terminal reductases. Knocking out *torA* increased the rate of PCA oxidation relative to the wildtype (Figure 7G and H). We found that the *dmsA* knockout strain had a PCA oxidation deficit during TMAO-driven PCA oxidation, which is consistent with the Dms complex promiscuously reducing many N-oxides (33,34) (Figure 7G and H). The *dmsAtorA* double knockout strain was faster at oxidizing PCA than the *dmsA* single knockout strain, implying that *C. portucalensis* MBL possesses other TMAO reductases (Figure 7I). We did not pursue a full study of the role of quinones during TMAO-driven PCA oxidation because we could not constrain the full activity.

### Nitrate-driven PCA oxidation provides a survival benefit for *C. portucalensis* MBL in a bioelectrochemical reactor

With a new understanding of the mechanism of biological PCA oxidation, we can assess its impact on bacterial fitness over longer time scales. After having grown to stationary phase in LB, *C. portucalensis* MBL continuously oxidizes PCA, so long as nitrate is available in a bioelectrochemical chamber with a working electrode poised to a reducing voltage (Figure 8). Upon spiking fresh nitrate into the reactor after PCA oxidation ceases, the oxidation immediately resumes, indicating that nitrate availability is the limiting factor for these cultures (Supplementary Figure 8.1). Adding more nitrate while PCA oxidation was active does not affect the current, which implies that the minimum threshold for the permissible nitrate concentration is relatively low (Supplementary Figure 8.1, green trace). After several days of oxidizing PCA with nitrate, the apparent rate of nitrate consumption goes down, as evidenced by a persistent current in the 24-hour spiking condition after the third day (Supplementary Figure 8.1, yellow trace). In the cultures that received addition nitrate, there was a slight but steady decrease in total current over time (Supplementary Figure 8.1, yellow and green traces). We did not assess whether there are analogous dynamics for PCA oxidation in the bioelectrochemical chamber when provided any the other terminal electron acceptors that we compared above.

**Figure 8.**
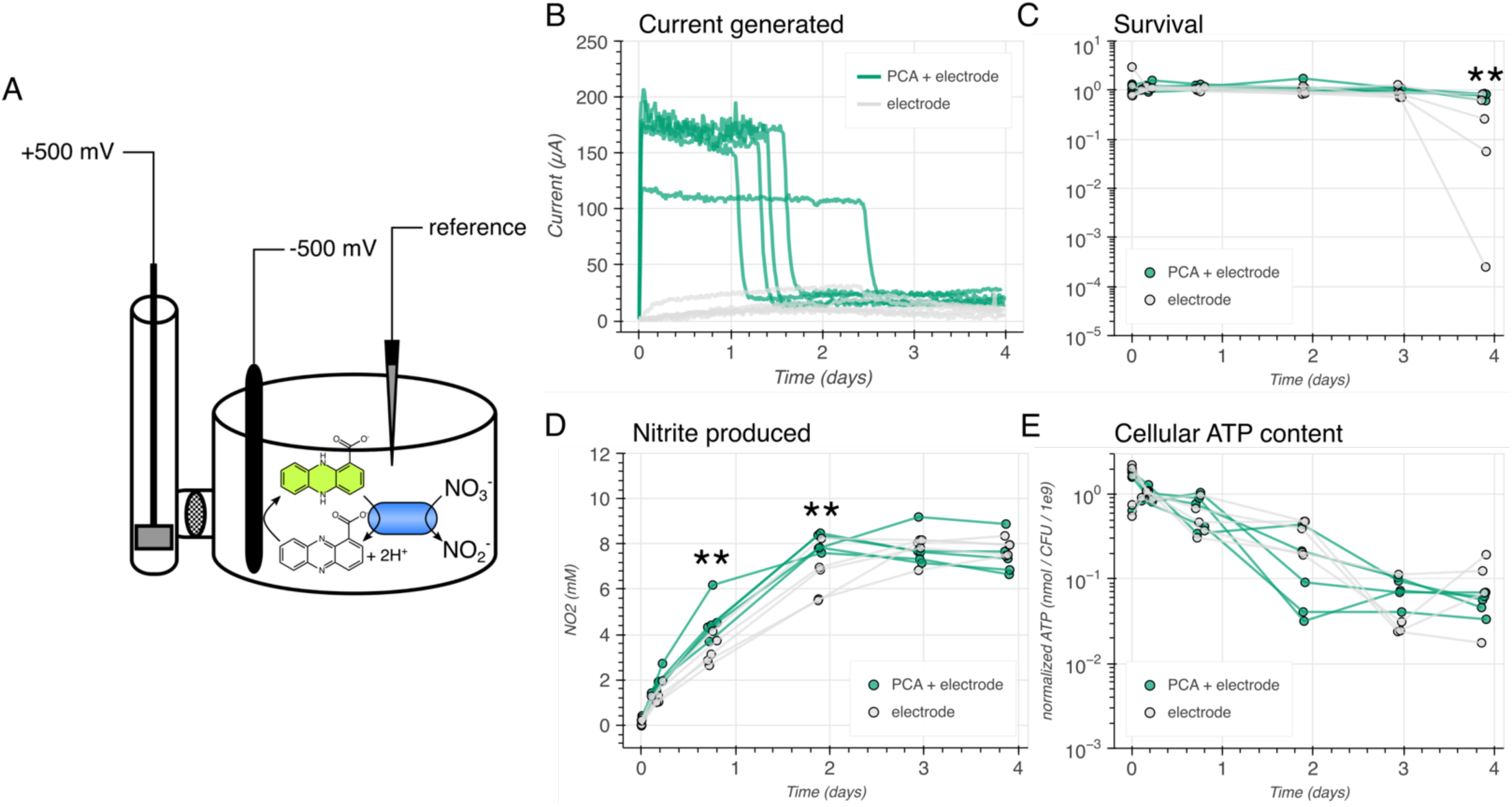
Physiological consequences of long-term PCA oxidation for C. portucalensis MBL. (A) Diagram of the bioelectrochemical reactor. The *C. portucalensis* MBL culture was incubated in the main chamber with a working electrode poised to -500 mV that continuously reduced any available PCA. The reference electrode communicated with the potentiostat to retain a constant voltage. Cells oxidized PCA when nitrate was present. In a sidearm, separated by a dense glass frit, the counter-electrode completed the circuit. Reduced PCA is green and oxidized PCA is colorless. (B) Current traces for biological replicates with and without provided PCA and an initial concentration of 10 mM nitrate. Samples were taken from these replicates at the times displayed in (C)-(E), where each data point corresponds to an independent biological replicate. (C) Timepoint measurements of survival in the culture, as determined by colony forming units (CFUs). The asterisks represent a Bonferroni-corrected statistically significant difference between the two conditions at the final timepoint (p = 0.0025). (D) The nitrite that was produced (from nitrate reduction) by the cultures over the experiment. At the third and fourth timepoints, there were Bonferroni-corrected statistically significant differences between the two conditions: p = 0.0029 and p = 0.0019, respectively. (E) The normalized ATP content per CFU over the time course. There were no statistically significant differences. The Bonferroni corrected threshold was p < 0.0083 given hypothesis testing across the six timepoints.

Having observed PCA oxidizing activity over multiple days, we compared three potential metrics of *C. portucalensis* MBL fitness (colony forming units (CFUs), cellular ATP content, and nitrite production) in two chamber bioelectrochemical reactor conditions, with and without PCA, both providing a reducing working electrode and an initial concentration of 10 mM nitrate as the terminal electron acceptor (Figure 8B-E). In the chamber with PCA, the cells converted nitrate to nitrite faster than in the chamber without PCA, which coincided with the loss of current as nitrite concentrations reached their asymptote (Figure 8B and D, third and fourth timepoints). Despite the PCA oxidation current ceasing between days one and three, the effect on survival was not apparent until day four, on which the cultures with PCA began to die (Figure 8C).

Throughout the time course, there were no samples that indicated a difference in cellular ATP content between the two conditions (Figure 8E). Although this experiment cannot speak to the underlying reason for differential cell survival, it is apparent that extended PCA oxidation was beneficial under these conditions, in which PCA was the sole electron donor provided.

## Discussion

In this work, we adapted an *E. coli* genetic toolkit to *C. portucalensis* MBL, a soil isolate. This enabled us to bring the study of the mechanism of biological PCA oxidation a step closer to its native soil environment. The observations that (1) any respirable terminal electron acceptor stimulates PCA oxidation and (2) that the presence of quinones is required for a large fraction of the oxidation rate imply a generalizable mechanism: any cell that can perform anaerobic respiration and has an accessible inner membrane should oxidize PCA. This is reflected in our inability to find a bacterium that does not oxidize PCA under suitable conditions (15,29).

Notably, had we not compared PCA oxidation by *C. portucalensis* MBL under differing pre-growth conditions, we may have had arrived at an errant model. For example, had we relied on oxic-pregrowth cultures, we would have concluded that the periplasmic nitrate reductase is not involved in PCA oxidation (Figure 4C).

When comparing how different organisms and different terminal electron acceptors induce PCA oxidation, it is important to keep in mind the biosynthetic capacity of the organism (both the presence of the pathway and its regulation) to make the various quinones (Supplementary Figure 2.1), as well as the ability of a given terminal reductase to use each of the quinones. For instance, *C. portucalensis* MBL synthesizes all three quinone described above, but the production of those quinones are differentially regulated depending on aerobic or anaerobic growth, and the periplasmic nitrate reductase only makes use of ubiquinone and menaquinone (not demethylmenaquinone) (Figure 2 and Supplementary Figure 2.1) (18,20,21). Meanwhile, *Pseudomonas aeruginosa* PA14 can produce *only* ubiquinone and has a very limited capacity for fermentative growth, which made it unfeasible to test quinone-null or double nitrate reductase mutants (24,40).

Species-specific quirks accounted for, the theoretical and experimental framework we describe in this report lends several testable predictions. First, it forecasts the genetic and chemical conditions under which other phenazines such as pyocyanin (PYO, E°′_1/2_ = -0.040 V) would be oxidized (Wang and Newman, 2008). PYO’s redox midpoint potential, being more positive than that of PCA, indicates that its oxidation could be mediated by ubiquinone or demethylmenaquinone but not menaquinone (Figure 2B). For the same reason, fumarate should not be a suitable terminal electron acceptor to drive PYO oxidation. A knockout of the gene *ubiE*, which disrupts ubiquinone and menaquinone synthesis but keeps demethylmenaquinone synthesis intact (Supplementary Figure 2.1), would enable a more detailed exploration of the interactions of phenazines, quinones, and the anaerobically respiring electron transport chain (Nitzschke and Bettenbrock, 2018; Sharma et al., 2012). While the *ubiCmenA* double knockout had a more severe PCA oxidation phenotype than the *menA* knockout alone, the *ubiC* single knockout phenotypes and the complementation results suggest that (demethyl-)menaquinones play a more significant role than ubiquinones (e.g., Figures 5A and 8A). This motivates further dissection of the role of specific quinones during PCA oxidation in future studies.

We observed that, in the absence of quinones, PCA oxidation can still be driven by terminal reductases in the case of nitrate (Figure 5A *menAubiCnapAnarZ-tlKO*, *menAubiCnarGnapA-tlKO*, and *menAubiCnarGnarZ-tlKO*), fumarate (Figure 8A *menAubiC-tlKO*), and DMSO (Figure 8D, *menAubiC-tlKO*). In other words, Category 1 PCA oxidation reactions (Table 1) play a subtle, but substantial role during anaerobic respiration. This may mean that either PCA is functionally substituting for a quinone, that there is an unknown other intermediate between PCA and the terminal reductase, or that there is an unappreciated abiotic oxidation of PCA in these contexts. The third case is unlikely: while nitrite that is produced during nitrate respiration can oxidize PCA at a low rate (Tsypin and Newman, 2021), its production requires that electrons be donated to the terminal nitrate reductase; without quinones, the next logical source of electrons would be the PCA, meaning that the PCA is ultimately oxidized by the terminal reductase and not nitrite. Furthermore, the fumarate and DMSO observations do not provide the same abiotic recourse as nitrate. There may well be an unknown factor that mediates electron transfer between PCA and the terminal nitrate and fumarate reductases. However, the existence of archaeal methanophenazine suggests that such a factor is not necessary. Methanophenazine is effectively the core phenazine molecule with a long aliphatic tail, and is the membrane electron carrier in *Methanosarcina mazei*, which lacks quinones (41). In other words, in *M. mazei*, we have an example of canonical electron shuttles being completely replaced by a phenazine in the membrane. Given this example and the fact that five different terminal reductase complexes catalyze PCA oxidation in the absence of quinones, the most parsimonious explanation is that reduced PCA replaces quinols as an electron donor to these complexes at an appreciable rate. The efficacy with which different phenazines may substitute for quinones likely depends on their hydrophobicity and diffusion within the plasma membrane.

The flexibility of conditions under which *C. portucalensis* MBL oxidizes PCA is underscored by the functional redundancy of the various quinones and of certain terminal reductases. This is most readily evident in the terminal nitrate reductases, where NarG, NarZ, and NapA each contribute different fractions of PCA oxidation depending on how the cultures were pre-grown. Assuming that the regulation of these genes is analogous to their orthologs’ regulation in *E. coli*, NarG and NapA are regulated primarily by Fnr in response to hypoxia and nitrate availability. In *E. coli*, NarZ is primarily regulated by RpoS in response to growth arrest. This regulatory schema corresponds to our observation that the *narZ* single knockout has the most severe phenotype in the slanted shaking pre-growth treatment (in which cultures reach stationary phase) and that the *narG* single knockout has the most severe phenotypes in the standing pre-growth treatment (Figure 4). This is further supported by our comparison of the *nap* and *nar* knockouts in *Pseudomonas aeruginosa*: *P. aeruginosa* has only one homolog of NarG that is regulated by Fnr, and its Nap operon is regulated by RpoS. Correspondingly, the *nap* knockout strain of *P. aeruginosa* has the more severe PCA oxidation deficit when pre-grown in shaking tubes to stationary phase, and the *nar* knockout strain has the more severe phenotype when pre-grown in non-shaking tubes. Thus, the ability of a cell to oxidize PCA is defined by both by its genetic content and regulatory state. The redundant pathways for PCA oxidation enable organisms like *C. portucalensis* MBL to perform the process under many different environmental conditions, all of which having in common electron flux through the electron transport chain and the quinone pool.

In the case of TMAO-driven PCA oxidation, we were unable to fully abolish the activity by knocking out terminal reductases. We found that the DMSO reductase, as represented by the catalytic subunit DmsA, contributed to PCA oxidation by TMAO (Figure 8G). This corresponds to the promiscuity of the DMSO reductase that has been reported in other organisms (34).

However, the *torA* knockout had no phenotype of on its own, and the *dmsAtorA* double knockout oxidized PCA faster than the *dmsA* knockout alone (Figure 8G). This implies that there is another TMAO reductase(s) in *C. portucalensis* MBL that also participates in TMAO-driven PCA oxidation. Indeed, *C. portucalensis* MBL has two homologs to the catalytic subunit of the *E. coli* TorYZ reductase (NCBI accession IDs NUH53377.1 and NUH54782.1), which may be the culprit (42). If knocking out these two genes in addition to *dmsA* and *torA* is not sufficient to abolish PCA oxidation activity with TMAO, the causative enzyme could potentially be identified via a mutant screen in the quadruple knockout genetic background, looking for loss of PCA oxidation activity during TMAO respiration.

In addition to illuminating the molecular mechanisms responsible for PCA oxidation, our work indicates that PCA oxidation may provide a fitness benefit for *C. portucalensis* MBL (Figure 12). We presume that such benefits will arise when the cells are starved for electron donors; in future follow-up studies, we predict that depletion of the internal carbon stores of the cells prior to a survival assay would be expected to magnify the effect. Given that PCA reduction can provide a fitness benefit for *Pseudomonas aeruginosa* (6,43), it may be possible to pair phenazine oxidizing and reducing bacteria, such that the complete PCA redox cycle supports the survival of both (44). This scheme would work best if the phenazine oxidizer has exclusive access to the terminal electron acceptor and that the phenazine reducer has exclusive access to the terminal electron donor (Figure 9A).

**Figure 9.**
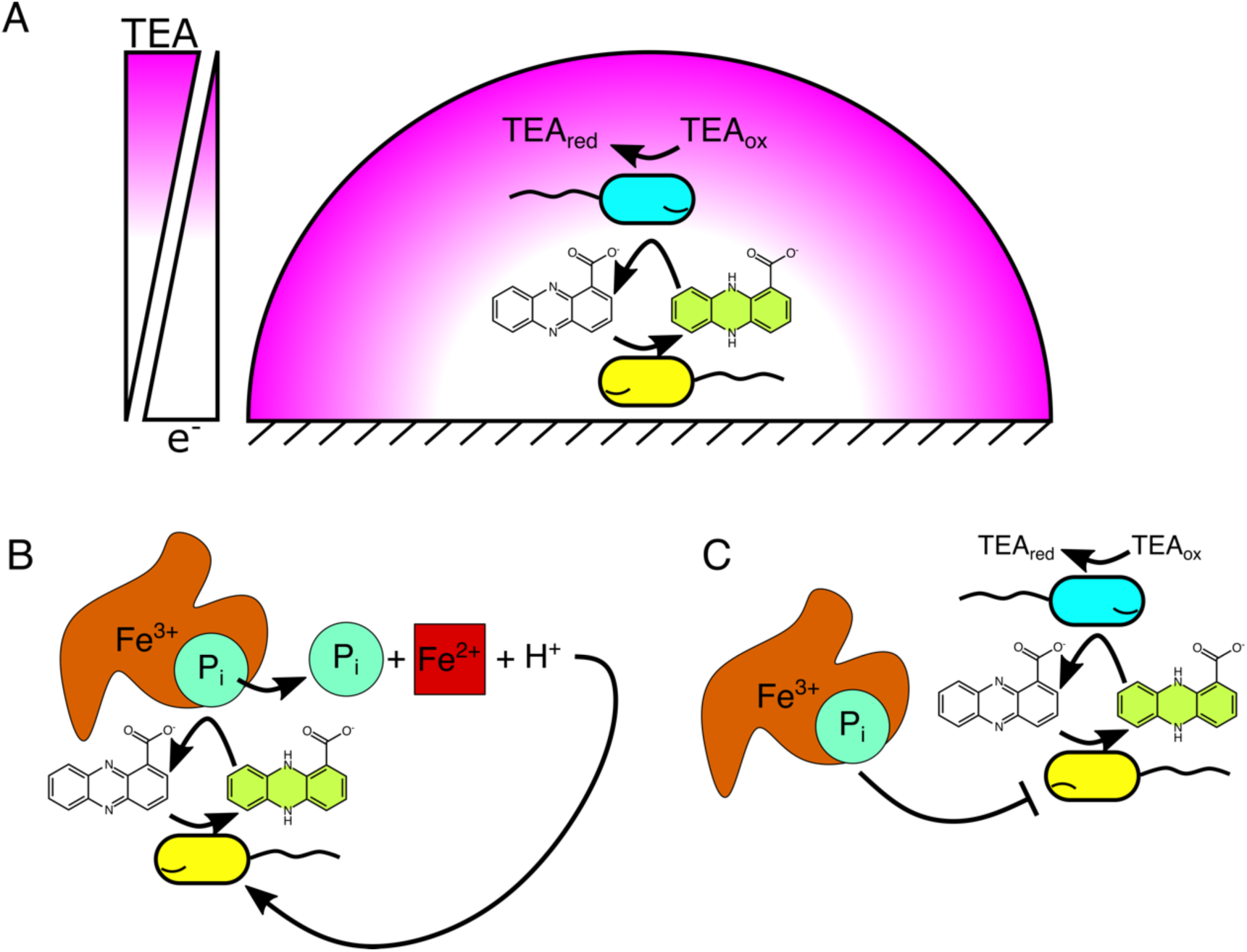
Model for how phenazine redox cycling may be mutualistic or competitive in microbial communities across redox gradients. (A) For a bacterium that finds itself starved for its preferred terminal electron acceptor (yellow cell under reducing stress), oxidized PCA can serve as an alternative. For a bacterium that finds itself starved for its preferred electron donor (cyan cell under oxidative stress), reduced PCA can serve as an alternative. Together, the two cells can more efficiently use bioavailable nutrients with the PCA redox cycle as a bridge. (B-C) For a bacterium that is relying on the abiotic oxidation of PCA to release insoluble nutrients (yellow cell, B), the presence of a PCA oxidizing species would starve it (cyan cell, C). This situation could occur if the environment contains a TEA that the blue cell can respire, but the yellow cell cannot. A specific example could pair a *P. aeruginosa* (as the yellow cell) and *C. portucalensis* MBL (as the cyan cell) in the presence of fumarate (31).

Beyond employing PCA oxidation to conserve energy, it is also possible for a bacterium to use PCA oxidation to compete against its neighbors (Figure 9B and C). Given that some bacteria appear to reduce PCA to release bioavailable iron and phosphorus (Figure 9B) (11,14), if an organism like *C. portucalensis* MBL were to intercept and oxidize the PCA before it reached the target mineral, the PCA reducer would remain starved for iron or phosphorus (Figure 9C). Thus, depending on the context, PCA oxidation may serve either mutualistic or competitive interactions between bacteria. Identifying these partnerships may enable new model systems for studying bacterial interactions beyond metabolite exchange or antibiotic secretion. Given the redox nature of these interactions, they are worthwhile to explore in bioelectrochemical reactors, where bacterial phenazine oxidation/reduction processes may compete with or supplement electrode activity. A clear direction for future studies is to assess the relative survival or fitness of different *C. portucalensis* MBL genotypes in electrode chambers. For example, a quinone null strain with only one of the three nitrate reductases subjected to a reducing electrode under anoxia would be forced to rely on electrons from PCA to drive nitrate reduction: will this culture survive better than one provided no PCA? No electrode? How might its PCA oxidation activity impact another organism grown in co-culture? The methods and results of this report open the field to ask detailed questions about the role of electron donors, separate from carbon sources, in bacterial metabolism.

We found that PCA oxidation is common and generalizable when it had not previously been observed or appreciated, which we were able to do by letting theory and comparative experiments be our guides. Biological PCA oxidation may play an important role in soil environments, inviting future exploration. We believe that the broader importance of this work is three-pronged. First, it gives an example of a conceptual framework for bridging thermodynamic reasoning, environmental microbiology, and cellular physiology. Second, it demonstrates of how mature our genetic engineering methods have become, allowing us to take an organism out of soil and develop a full genetic toolkit to test hypotheses in the span of a few years (15,16). Third, it underscores the importance of testing phenotypes across multiple physiological conditions, as some gene activities may be cryptic under a given gene regulatory mode. There are likely many other similar redox metabolisms in the environment that await discovery, and our report provides an example of how they may be pursued.

## Materials and Methods

### Statistical comparisons

All data and code for analysis are available on GitHub (https://github.com/ltsypin/Cportucalensis_genetic_mech) and via the Caltech library (https://doi.org/10.22002/tdng7-twd27). Given data that are not necessarily normally distributed, we employed bootstrapping for all our hypothesis testing, which we implemented as a two-tailed variant of the empirical t-statistic described by Efron and Tibshirani (45) (Algorithm 16.2 in the book). Specifically, when comparing two sets of datapoints, we tested the null hypothesis that they possessed the same means. We did this via the following algorithm:

1) Suppose you have two data sets X = {x_1_, x_2_, …, x_n_} and Y = {y_1_, y_2_, …, y_m_}.
2) If both sets have at least three elements, calculate the observed difference between their means:

a. µ_x_ = 1/n Σ(X)
b. µ_y_ = 1/m Σ(Y)
c. d_obs_ = µ_x_ - µ_y_.
3) The observed t-statistic, t_obs_ = d_obs_ / √(σ_X_^2^/n + σ_Y_^2^/m), where σ_X_^2^ is the variance of X and σ_Y2_ is the variance of Y.
4) In order to generate the distribution that simulates the null hypothesis, shift the means of both sets to be equal:

a. µ_tot_ = (Σ(X) + Σ(Y)) / (n + m)
b. X’ = X - µ_x_ + µ_tot_
c. Y’ = Y - µ_y_ + µ_tot_.
5) Generate the null hypothesis distribution by drawing 1,000,000 bootstrap replicates with replacement from the shifted data sets. For each replicate

a. Draw a size-n sample with replacement from X’: x’
b. Draw a size-m sample with replacement from Y’: y’
c. d_b_ = 1/n Σ(x’) – 1/m Σ(y’)
d. t_b_ = d_b_ / √(σ_x’_^2^/n + σ_y’_^2^/m), where σ_x’_^2^ is the variance of x’ and σ_y’_^2^ is the variance of y’.
6) Calculate the p-value of t_obs_ relative to the distribution of T = {t_1_, t_2_, … t_1,000,000_}:

a. Count z, the number of elements of D, where |t_j_| ≥ |t_obs_| (the absolute values are what make this a two-tailed test)
b. p = z/1,000,000.
7) For determining significance, apply the Bonferroni correction by dividing the standard threshold p-value (p = 0.05) by the number of concurrent pairwise comparisons. E.g., if there are 10 strains compared pairwise, statistical significance is at p < 0.005.

### Reagents

All chemical compounds used were purchased from Sigma-Aldrich. We want to note for future researchers that we encountered batch effects with different lots of TMAO: some lots (particularly the hydrated TMAO stocks) abiotically oxidize PCA while others do not. With TMAO, it may be important to try a couple batches of the compound.

### Strains and culturing

We employed wildtype and mutant *C. portucalensis* MBL and *P. aeruginosa* UCBPP-PA14 in this study (Table 3). How the mutants were generated is described in the following section and in the Supplement. In brief, all strains were cultured in lysogeny broth (LB) prior to PCA oxidation assays. Each pre-growth in 5 mL LB was for 17 hours at 30 °C in borosilicate glass culture tubes. In Figures 4 and 6, we compare pre-growth in slanted shaking tubes (250 rpm) and standing tubes sealed with parafilm to reduce oxygen permeance. In the bioelectrochemical studies (Figure 8 and Supplementary Figure 8.1), wildtype *C. portucalensis* MBL was first pre-grown overnight in shaking slanted tubes and then outgrown to a larger volume before concentrating the cells and inoculating them into the reactors (see Supplement for a detailed protocol). For all other experiments, *C. portucalensis* MBL strains were pre-grown in standing, sealed tubes without supplemented nitrate.

**Table 3.**
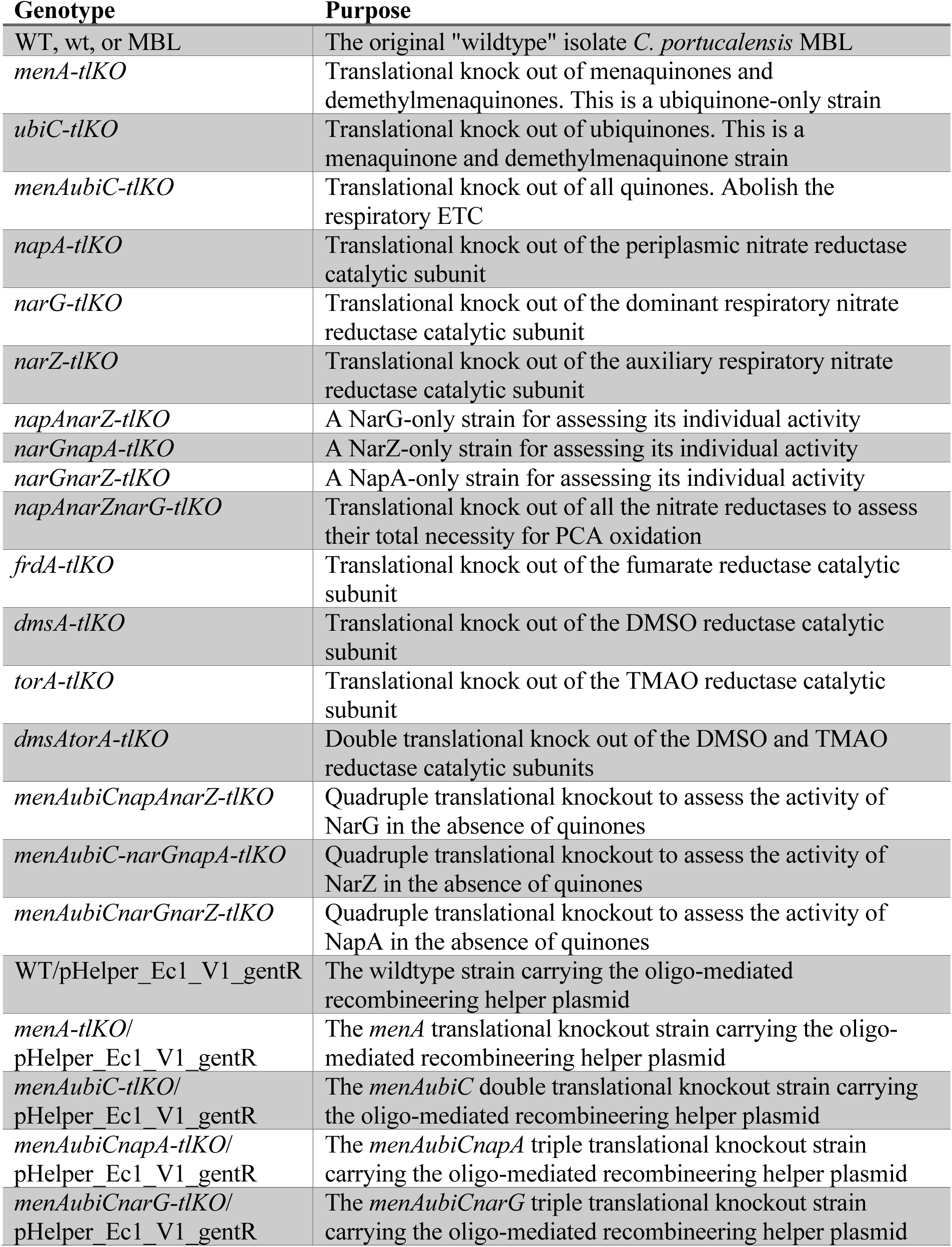
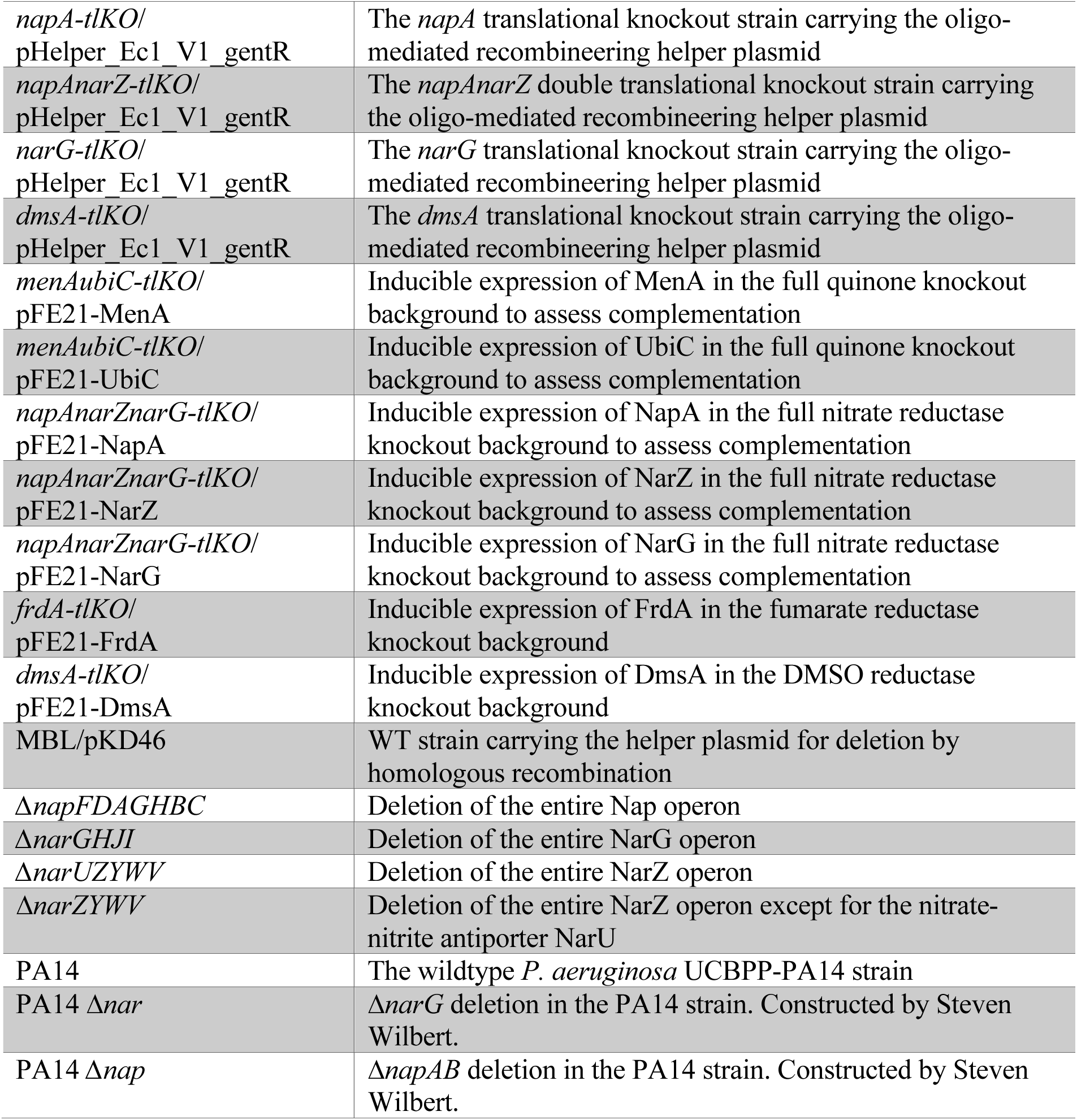
Strains used in this study.

### PCA oxidation assays

The basal PCA oxidation assay medium was composed of 20 mM potassium phosphate buffer (pH 7-7.1); 1 mM sodium sulfate; 10 mM ammonium chloride; and 1× freshwater salt solution (17.1 mM sodium chloride, 1.97 mM magnesium chloride, 0.68 mM calcium chloride, and 6.71 mM potassium chloride). Reduced PCA was prepared in this basal medium at a concentration of 1.2 mM by electrolysis (working electrode poised to -500 mV). Reduced PCA was the only electron donor added to the assay medium. We performed PCA oxidation assays either by tracking the fluorescence of reduced PCA over time in a plate reader (BioTek Synergy 4 or HTX) housed in a Coy-brand anoxic chamber or by tracking the current generated by a culture when provided PCA and an electrode poised to continuously reduce any PCA that the cells oxidized. The current traces were measured using a Gamry potentiostat, and these experiments were conducted in an mBraun-brand anoxic chamber, which scrubs hydrogen from the headspace. A detailed protocol for the plate reader assay is available via protocols.io (dx.doi.org/10.17504/protocols.io.bp2l6xm6dlqe/v1), and an explanation of how our electrode chamber assay differed from a previously published protocol (48) is available in the Supplement.

### Genetic engineering of C. portucalensis MBL

Detailed protocols for genetically engineering *C. portucalensis* MBL are available in the supplement. In brief, we employed both λRed-mediated homologous recombination to delete endogenous operons (26) and oligo-mediated recombineering to cause translational knockouts of genes of interest (27). The oligonucleotides and primers we used to accomplish this are listed in Tables 4 and 5. For generating the deletion strains, we used the helper plasmids pKD46 (for the λRed machinery), pKD4 (for the kanamycin resistance cassette flanked by FRT sites), and pCP20 (for the FLPase). For generating the translational knockouts, we used the helper plasmid pHelper_Ec1_V1_gentR, which was derived from pORTMAGE-Ec1 (27,46) by adding a SacB gene for sucrose counterselection to cure the plasmid. These helper plasmids are listed in Table 6. We transformed *C. portucalensis* MBL by electroporation: 200 Ω, 25 µF, 2.5 kV in 2 mm gap cuvettes. Our protocols for preparing electrocompetent *C. portucalensis* MBL, the Datsenko-Wanner knockouts, and the oligo-mediated recombineering are available via protocols.io:

1. Electrocompetent cells: dx.doi.org/10.17504/protocols.io.kqdg3x7r7g25/v1
2. Datsenko-Wanner knockouts: dx.doi.org/10.17504/protocols.io.ewov1q5e7gr2/v1
3. Oligo-mediated recombineering: dx.doi.org/10.17504/protocols.io.eq2lyj1rwlx9/v1

**Table 4.**
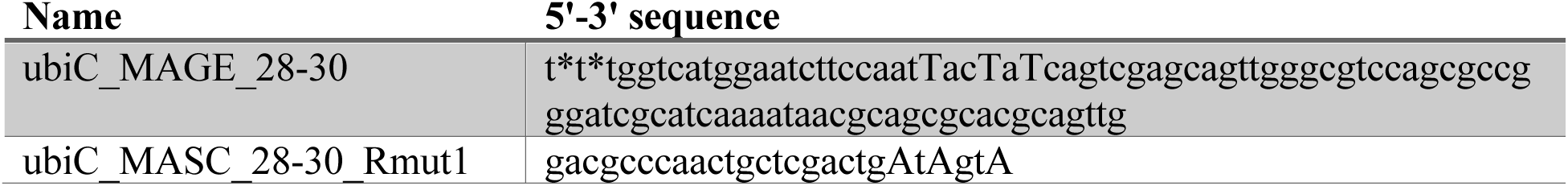
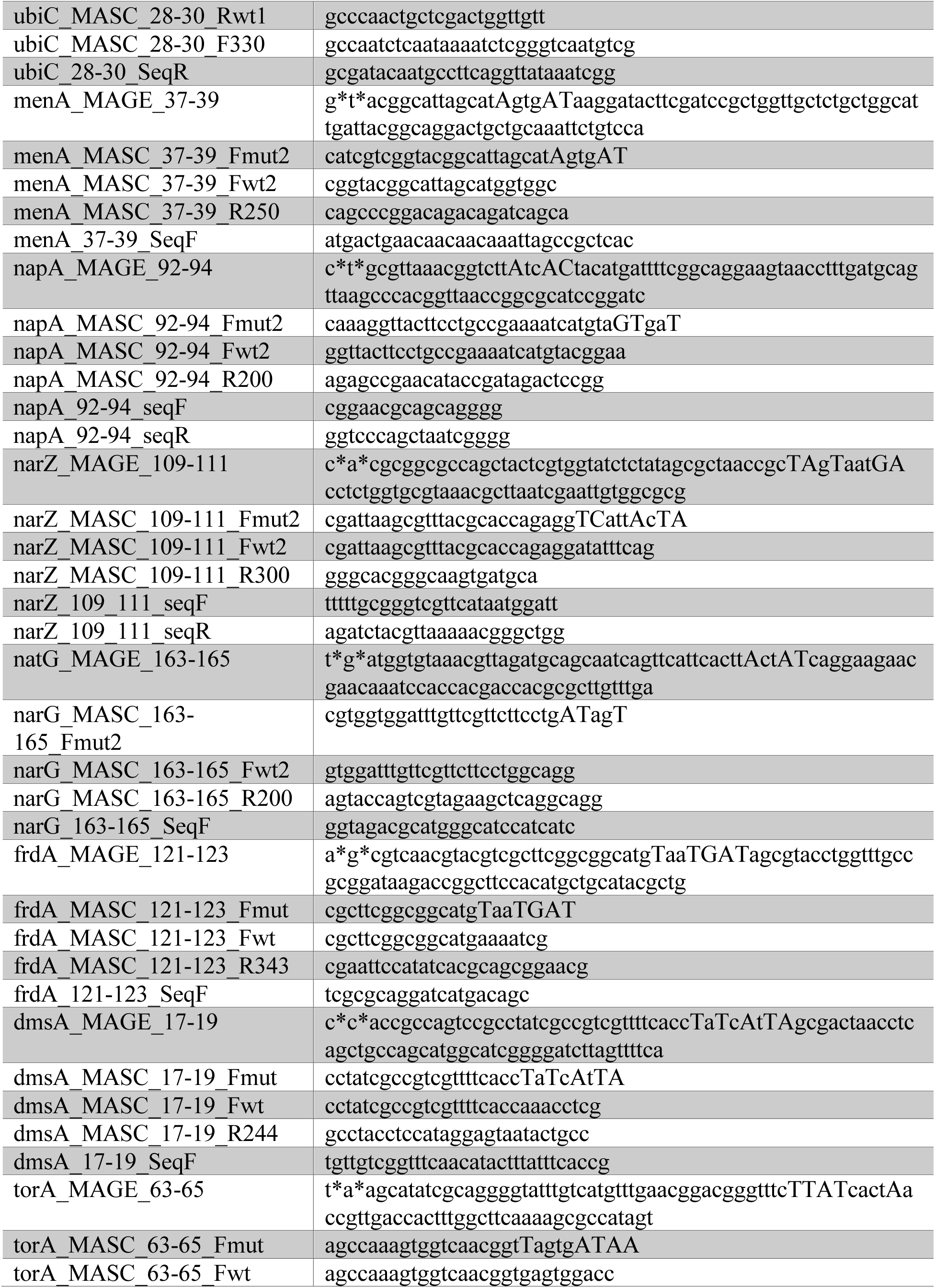
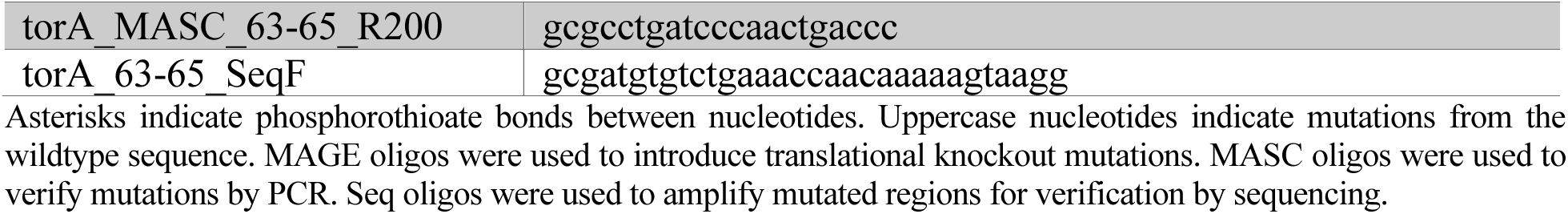
Oligos for translational knockouts.

**Table 5.**
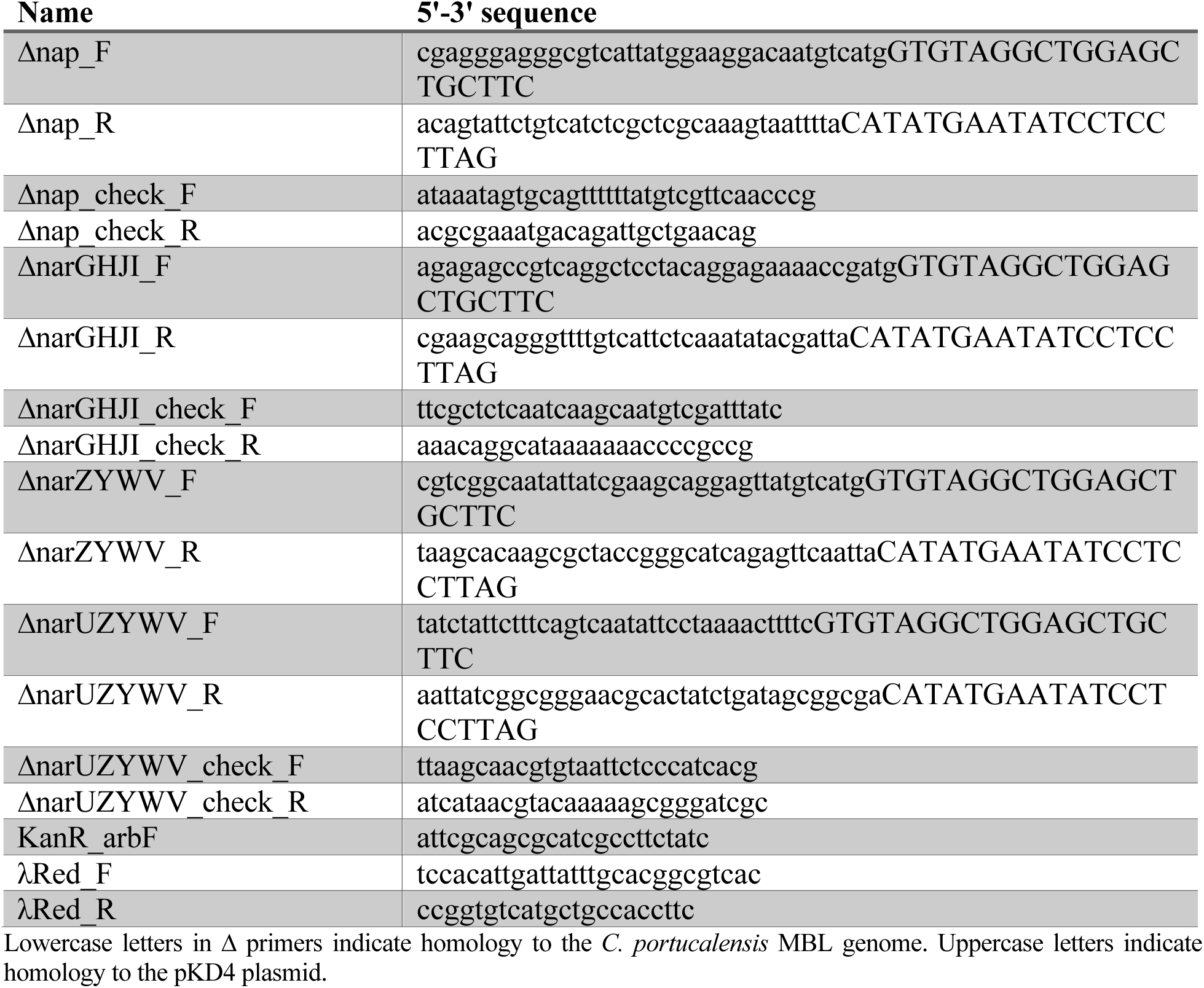
Oligos for Datsenko-Wanner deletions.

**Table 6.** Plasmids employed in the study.

| Name | Purpose | Reference |
| --- | --- | --- |
| pKD46 | λRed helper, temperature-sensitive origin of replication, confers ampicillin resistance | (26) |
| pKD4 | Kanamycin vector with FRT sites | (26) |
| pCP20 | FLPase vector, temperature-sensitive origin of replication, confers ampicillin resistance | (26) |
| pHelper_Ec1_V1_gentR | Oligo recombineering helper vector, cloned into backbone with SacB, confers gentamicin resistance | Scott Saunders, adapted from (27) |
| pFE21-sfGFP | Tetracycline-inducible promoter driving gene expression, confers kanamycin resistance | (47) |
| pFE21-MenA | pFE21 for inducible MenA expression | This work |
| pFE21-UbiC | pFE21 for inducible UbiC expression | This work |
| pFE21-NapA | pFE21 for inducible NapA expression | This work |
| pFE21-NarZ | pFE21 for inducible NarZ expression | This work |
| pFE21-NarG | pFE21 for inducible NarG expression | This work |
| pFE21-FrdA | pFE21 for inducible NarG expression | This work |
| pFE21-DmsA | pFE21 for inducible DmsA expression | This work |

### Cloning and induction of complementation vectors

We cloned the complementation vectors using the plasmid pFE21 as a backbone for inducible expression of wildtype *C. portucalensis* MBL genes (47). We used Gibson assembly to construct the new vectors from the pFE21 backbone and gene amplicons, and the primers we used are listed in Table 7. We induced the vector by incubating the cultures with 50 nM anhydrotetracycline during the pre-growth phase of the assay. The 50 nM concentration induction was deduced from an assay with pFE21-sfGFP, measuring GFP fluorescence on a BioTek Synergy 4 plate reader during aerobic growth (data not shown).

**Table 7.**
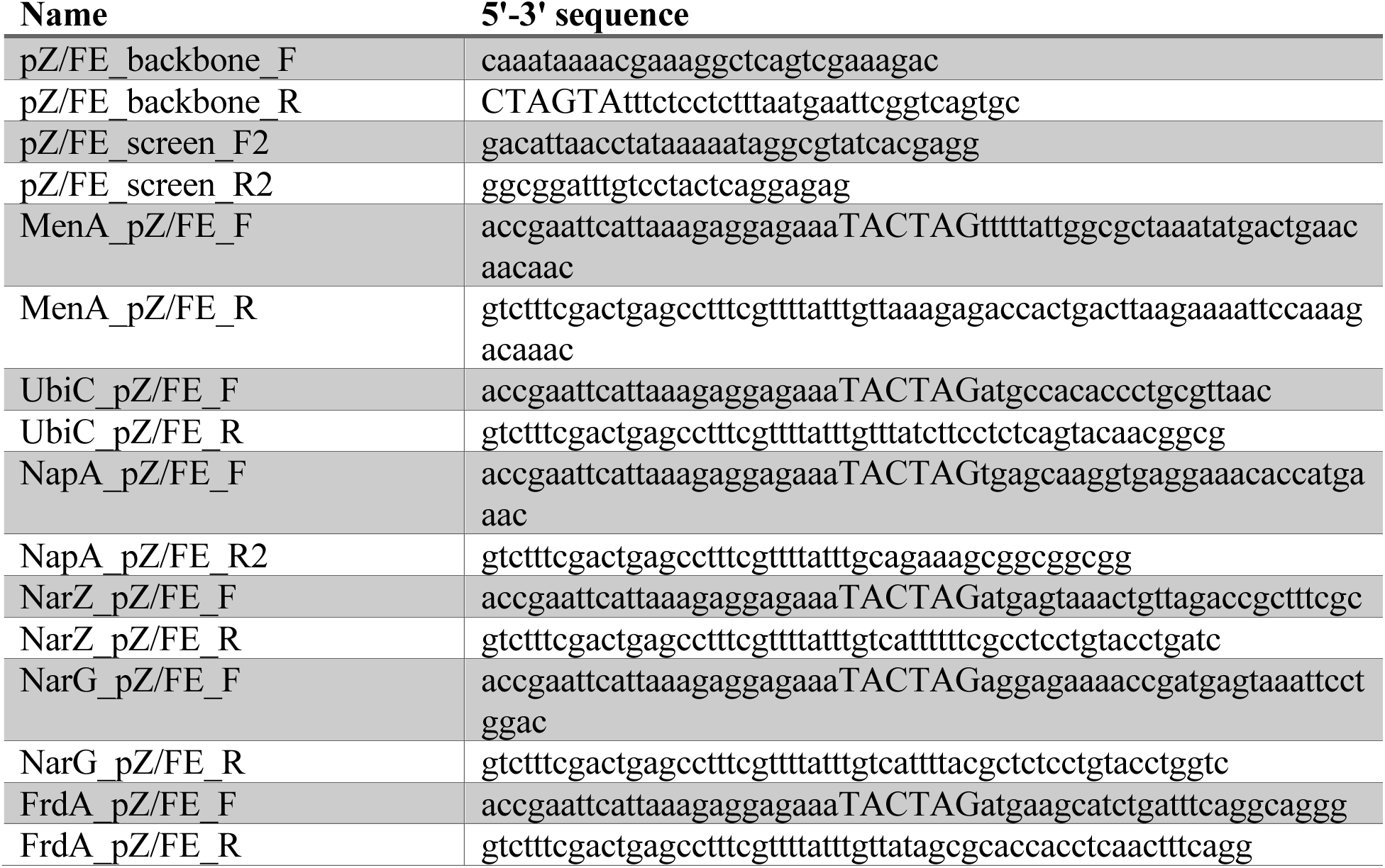
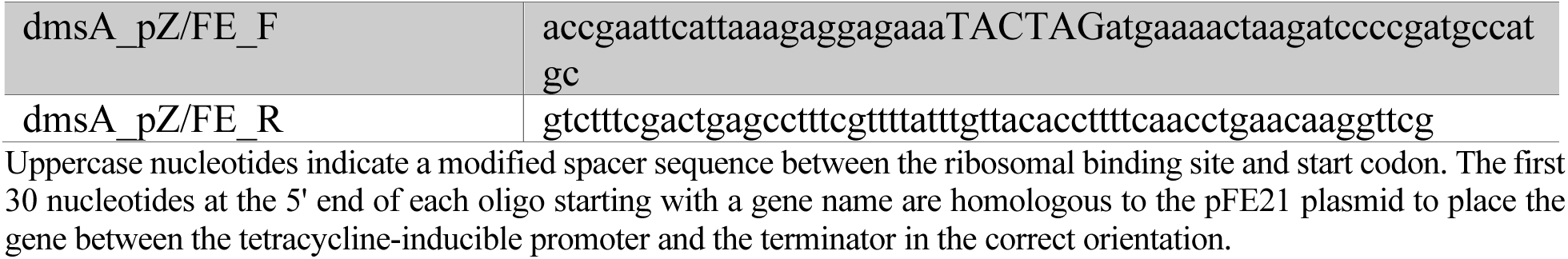
Oligos for building complementation vectors.

## Acknowledgements

We thank Steven Wilbert, John Ciemniecki, Chelsey VanDrisse, Georgia Squyres, Avi Flamholz, and Julian Wagner for helpful technical feedback and general support throughout this work. We thank Maxim Tsypin for pointing us to Efron and Tibshirani’s implementation of the bootstrapped hypothesis test. LMZT was supported by an NSF graduate research fellowship, and additional resources used in the study came from grants to DKN from the NIH (1R01AI127850-01A1) and Doren Family Foundation.

## Author contributions (CRedIT Taxonomy)

**LMZT**: conceptualization, data curation, formal analysis, funding acquisition, investigation, methodology, project administration, software, supervision, validation, visualization, writing – original draft preparation, writing – review and editing.

**SHS**: methodology, supervision, writing – review and editing.

**AWC**: investigation, writing – review and editing.

**DKN**: conceptualization, funding acquisition, methodology, project administration, resources, supervision, writing – review and editing.

## Abbreviations

(PCA): phenazine-1-carboxylic acid
(ETC): electron transport chain
(TEA): terminal electron acceptor
(NO_3-_): nitrate
(NO_2-_): nitrite
(N_2_O): nitrous oxide
(NO): nitric oxide
(Fum^2-^): fumarate
(Succ^2-^): succinate
(DMSO): dimethyl sulfoxide
(DMS): dimethyl sulfide
(TMAO): trimethylamine-N-oxide
(TMA): trimethylamine
(UQ): ubiquinone
(UQH_2_): ubiquinol
(MQ): menaquinone
(MQH_2_): menaquinol
(DMQ): demethylmenaquinone
(DMQH_2_): demethylmenaquinol

---

**Supplementary Figure 2.1.**
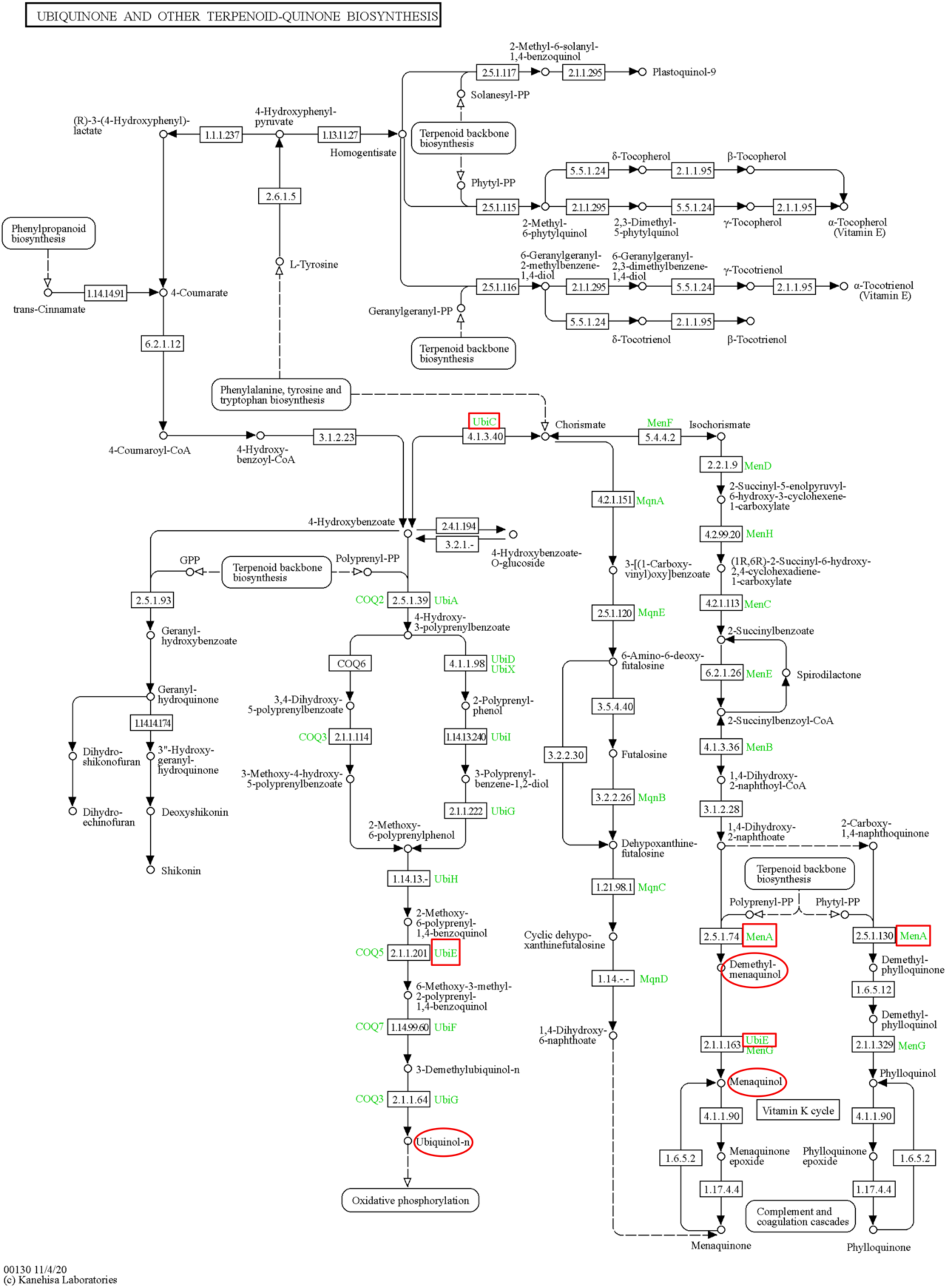
KEGG reference pathway for quinone biosynthesis (map00130). The relevant genes are indicated by red boxes and the relevant quinones by red ellipses. Note: *MenG* is a homolog to *UbiE* from photosynthetic organisms and is not present in γ-Proteobacteria like *C. portucalensis* MBL; the alternative pathway for menaquinone biosynthesis via futalosine (the *Mqn* genes) is also absent in *C. portucalensis* MBL (16,22,23). Loss of *UbiC* results in the loss of ubiquinones. Loss of *MenA* results in the loss of menaquinones and demethylmenaquinones. Loss of *UbiE* results in the loss of ubiquinones and demethylmenaquinones. Note: *Pseudomonas aeruginosa* only has ubiquinones in its ETC under both aerobic and anaerobic growth conditions (24). Pathway diagram used with permission from Kanehisa Laboratories (permission received November 6, 2023) (25).

**Supplementary Figure 4.1.**
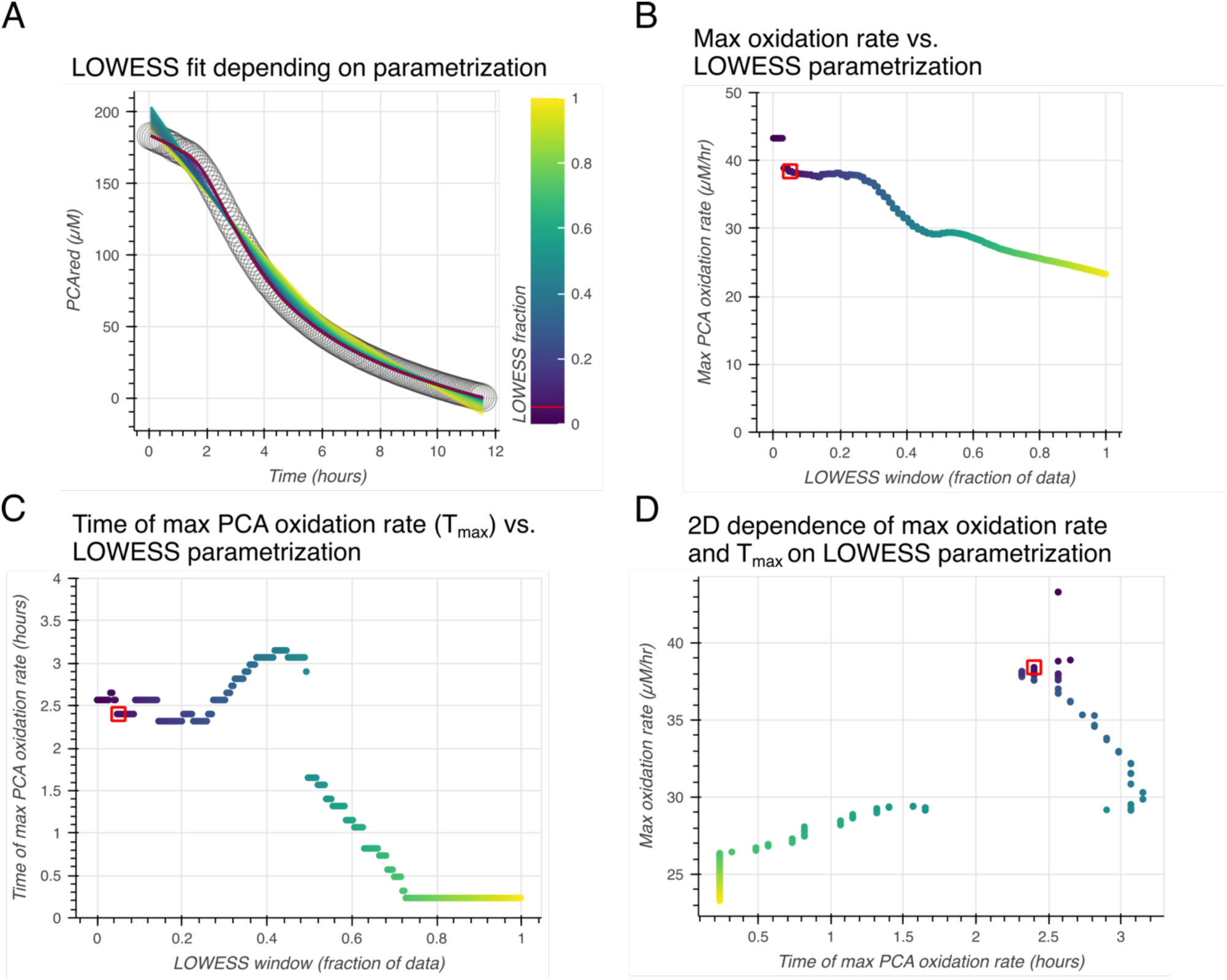
Quality of LOWESS fit depends on parametrization of its scanning window. The output of the LOWESS algorithm depends on one key parameter: the fraction of data that it considers for each smoothing window. (A) Examples of different fits to the same data as in Figure 4A, scanning the fraction parameter from 0 to 1. The value that was used for the analysis, 0.05, is in red. (B) How the estimated maximum PCA oxidation rate depends on the scanning window. The red square indicates the value at 0.05. (C) How the time to half of PCA being oxidized depends on the scanning window. The red square indicates the value at 0.05. (D) A 2D representation of both the maximum oxidation rate and the time at which it occurs. The red square indicates the value at 0.05. 0.05 was chosen as the window for all the analyses in this report as it appeared to give stable outputs while using a minimal window for fitting.

**Supplementary Figure 4.2.**
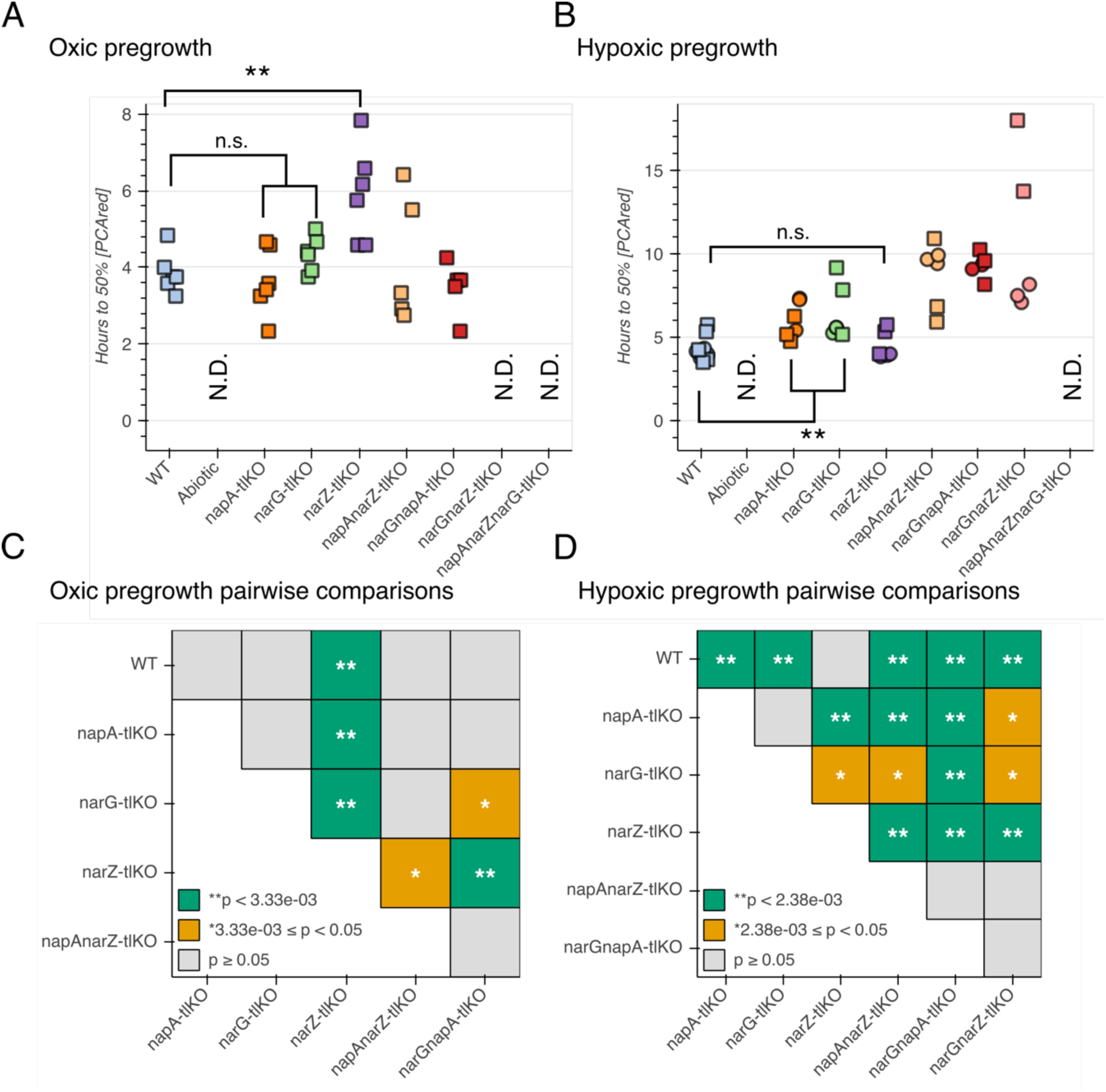
Nitrate reductase phenotypes as measured by time to oxidize half of provided PCA. (A) The time it took the assayed strains to oxidize half of the provided PCA after oxic pregrowth. (B) The time it took for the assayed strains to oxidize half of the provided PCA after hypoxic pregrowth. (C) Pairwise comparisons of the mean times required to oxidize 50% of the provided PCA. Given 15 comparisons, the Bonferroni-corrected p-value threshold for significance is p < 0.00333. (D) Likewise, pairwise comparisons of the strains after hypoxic pregrowth. Given 21 comparisons, the Bonferroni-corrected p-value threshold for significance is p < 0.00238. N.D. (not detected): strains that did not oxidize half of the provided PCA over the course of the assay (48 hours). These results support the same conclusions as the ones in Figure 4: after oxic pregrowth, NarZ is the dominant contributor to PCA oxidation; after hypoxic pregrowth, all three nitrate reductases contribute to PCA oxidation, but NarG is dominant.

**Supplementary Figure 4.3.**
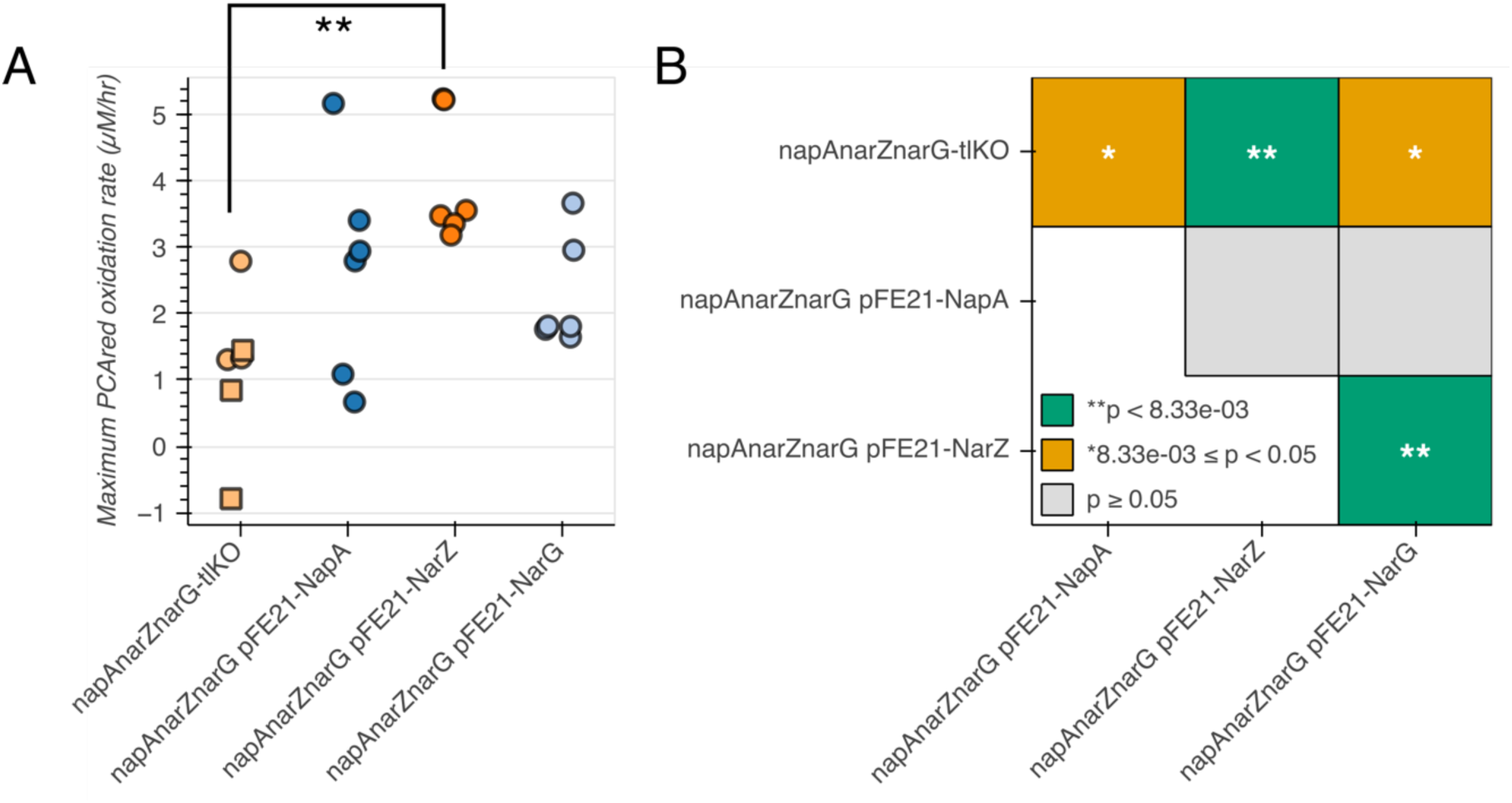
Complementation of nitrate reductase knockouts after overnight standing pre-growth. (A) The maximum PCA oxidation rate for the triple knockout strain and overexpressed individual nitrate reductases in that genetic background. Squares represent the means of technical triplicates and circles represent independent biological replicates. The negative value in the triple knockout background indicates that the cells were further reducing the provided stock of PCA, rather than oxidizing it. (B) Pairwise statistical tests against the null hypothesis that there is no difference between the mean maximum oxidation rates of two given genotypes. Given six comparisons, the Bonferroni-corrected p-value threshold for significance is p < 0.00833.

**Supplementary Figure 5.1.**
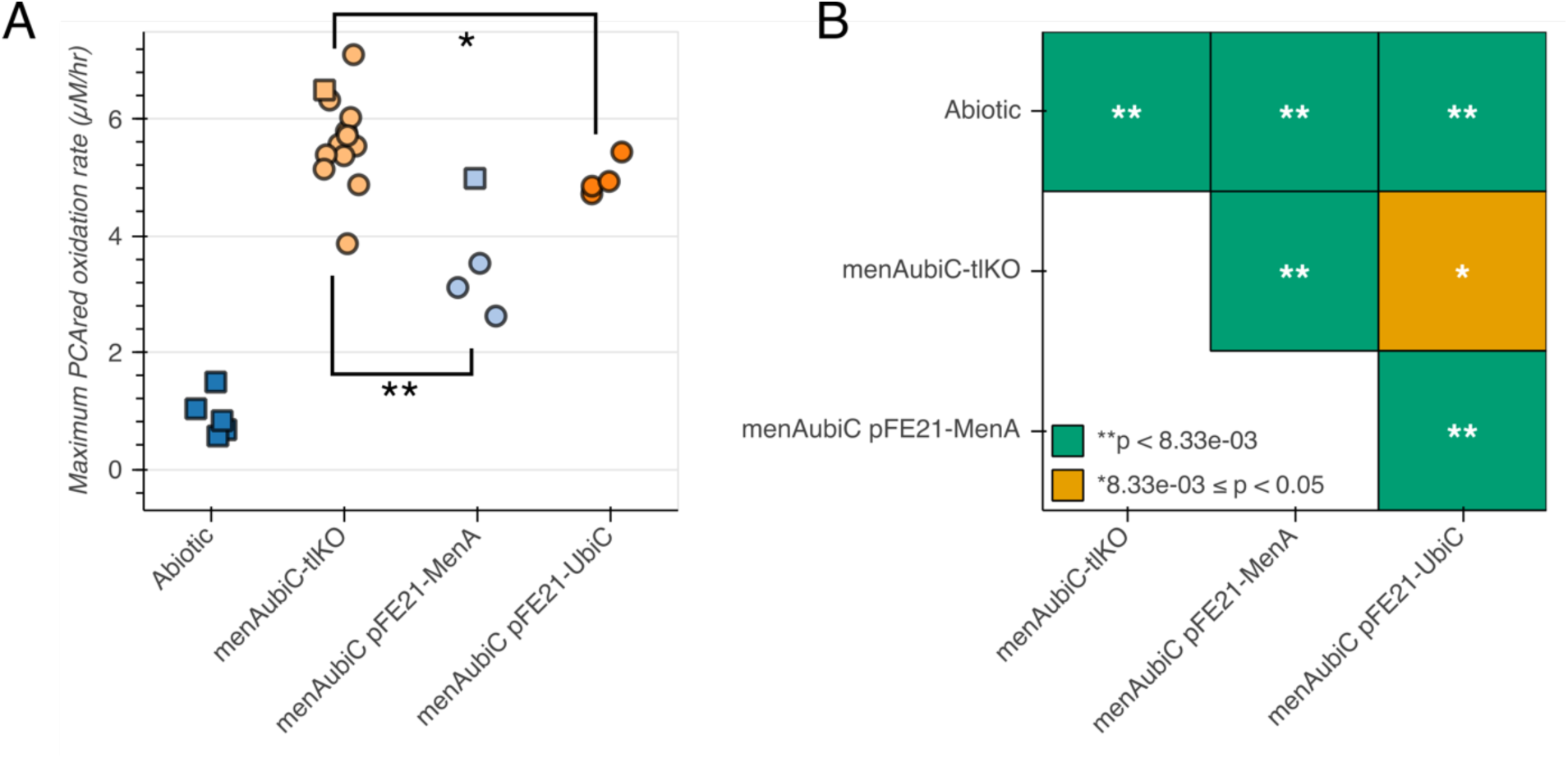
Overexpression of MenA and UbiC does not complement the PCA oxidation phenotype of the menAubiC double mutant. (A) Maximum PCA oxidation rates in complemented quinone knockout backgrounds. (B) Pairwise comparisons of the mean maximum PCA oxidation rates in (C). Given six comparisons, the Bonferroni-corrected threshold for significance is p < 0.00833.

**Supplementary Figure 8.1.**
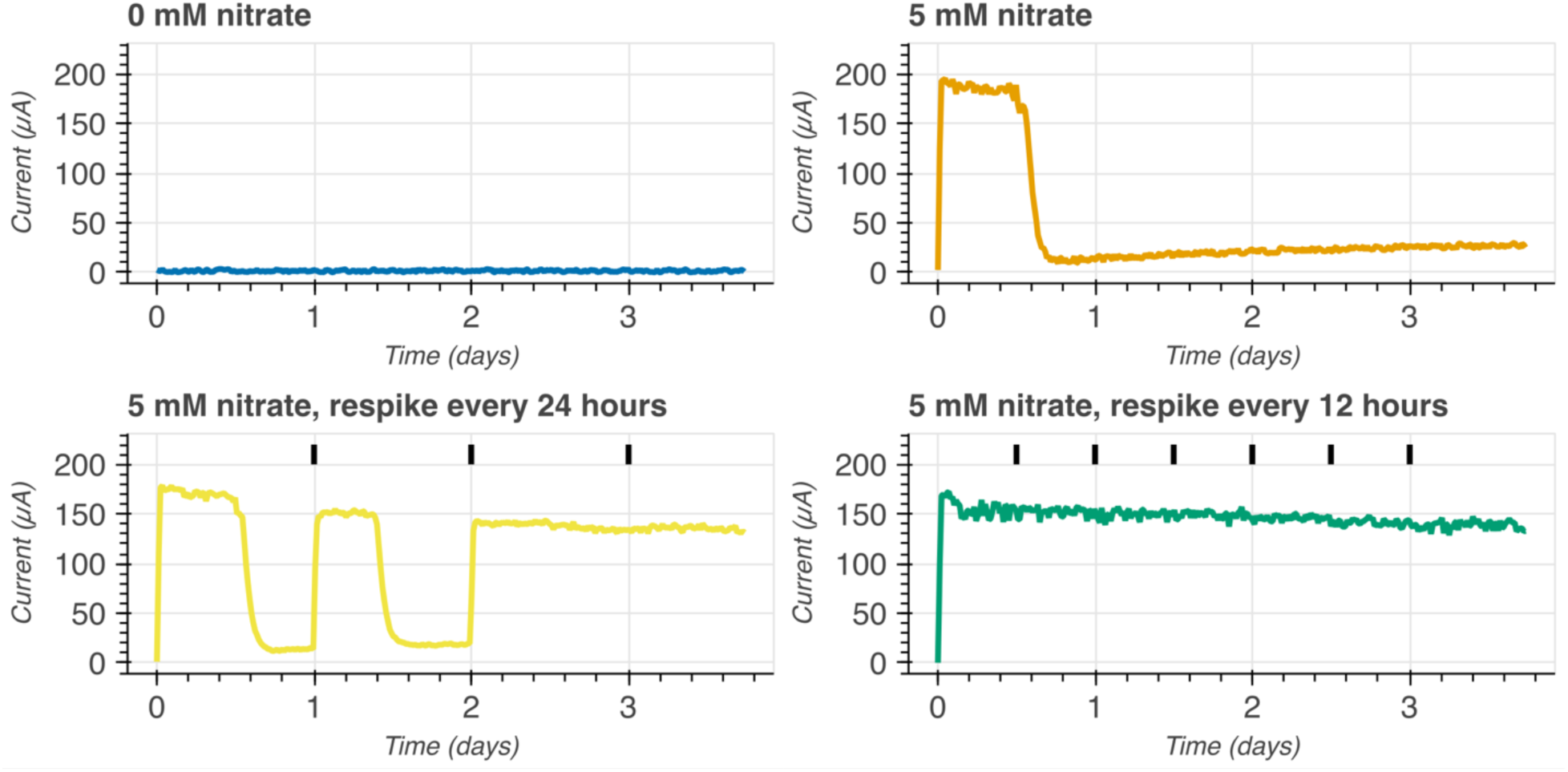
PCA oxidation depends on nitrate availability in a bioelectrochemical reactor. Each chart presents a time course of current (µA on a linear scale) during incubation of *C. portucalensis* MBL in a bioelectrochemical reactor with a working electrode that continuously reduces PCA. Current indicates that the culture is oxidizing PCA. Vertical black bars represent the timing of nitrate spiking, when appropriate. Each chart is titled according to the initial concentration of nitrate in the medium and the spiking schedule.

## Supplement

### Electrode chamber assay for PCA oxidation

The electrode chamber assays were performed as described in Suzanne Kern’s PhD thesis (48), Chapter 7, with the following modifications:

1. The medium for the assay was the phenazine oxidizer basal medium, as described in the main text.
2. The glass components of the electrode chambers were extensively cleaned prior to each experiment: they were soaked in methanol for at least 2 hours, soaked in a potassium hydroxide solution overnight (added a couple drops of potassium hydroxide to chambers filled with lab-pure water), soaked in 10% hydrochloric acid overnight, and then muffled at 980 °F for four hours, prior to assembly with stirbars and o-rings and autoclaving.
3. The polarity of the working and counter electrodes was flipped, with the working electrode poised to -500 mV relative to the reference electrode.
4. The graphite working electrodes (Alfa Aezar Cat. No. 14738) were polished with a Kim Wipe until shiny prior to soaking in 70% ethanol to sterilize them prior to chamber assembly.
5. The custom platinum mesh counter electrodes were also soaked in 70% ethanol to be sterilized prior to chamber assembly.
6. The reference electrodes were BASi Cat. No. MW-2030 instead of BASi Cat. No. RE-5B.
7. Instead of directly submerging the reference electrodes into the electrode chamber, they were inserted into ultrapure agarose (1% agarose in 3 M NaCl) in a 200 µL gel-loading pipette tip to create a stable salt bridge (also dipped into 70% ethanol to sterilize prior to use). The electrode-pipette tip connection was secured with Parafilm after the agarose had cured.
8. *C. portucalensis* MBL was pre-grown for these experiments as follows:

a. Grow overnight 5 mL LB cultures at 30 °C, shaking slanted tubes at 250 rpm.
b. Measure OD_600_ of overnight cultures and inoculate 250 mL LB in a 1 L Erlenmeyer flask to an OD_600_ = ∼0.06.
c. Grow the 250 mL LB cultures, shaking at 250 rpm at 30 °C, until they reach OC_600_ =∼2.8 (approximately 4-6 hours).
d. Pellet all 250 mL in one bottle by spinning for 10 minutes at 6000 × g at room temperature.
e. Resuspend cell pellet in 25 mL of the medium used for electrode chamber experiments and transfer to 50 mL conical tube.
f. Twice more, pellet the cells by spinning for 8 minutes at 5500 × g at room temperature.
g. After the second spin, resuspend in 6 mL of the experiment medium
h. Measure OD_600_ and adjust to a final value of OD_600_ = 75.
i. Bring cell suspension into mBraun anoxic chamber
j. Inoculate 1 mL into each appropriate electrode chamber.
9. ATP and CFUs were measured as described in Glasser et al. (6)

